# Combined neural tuning in human ventral temporal cortex resolves the perceptual ambiguity of morphed 2-D images

**DOI:** 10.1101/460337

**Authors:** Mona Rosenke, Nicolas Davidenko, Kalanit Grill-Spector, Kevin S. Weiner

## Abstract

We have an amazing ability to categorize objects in the world around us. Nevertheless, how cortical regions in human ventral temporal cortex (VTC), which is critical for categorization, support this behavioral ability, is largely unknown. Here, we examined the relationship between neural responses and behavioral performance during the categorization of morphed silhouettes of faces and hands, which are animate categories processed in cortically adjacent regions in VTC. Our results reveal that the combination of neural responses from VTC face- and body-selective regions more accurately explains behavioral categorization than neural responses from either region alone. Furthermore, we built a model that predicts a person’s behavioral performance using estimated parameters of brain-behavioral relationships from a different group of people. We further show that this brain-behavioral model generalizes to adjacent face- and body-selective regions in lateral occipito-temporal cortex. Thus, while face- and body-selective regions are located within functionally-distinct domain-specific networks, cortically adjacent regions from both networks likely integrate neural responses to resolve competing and perceptually ambiguous information from both categories.

---

We categorize objects within our visual world countless times each day and within a split second (Grill-Spector and Kanwisher 2005; Thorpe, Fize, and Marlot 1996). Despite this frequent occurrence and undisputable relevance to everyday life, it is presently unknown how regions and networks within the human brain support efficient and accurate visual categorization, especially for ambiguous stimuli (Hasson et al. 2001; Andrews et al. 2002; Moutoussis and Zeki 2002). Prior research has shown that human ventral temporal cortex (VTC) is critical for visual categorization (Haxby et al. 2000; Grill-Spector and Weiner 2014) and that shape is a key feature for neural categorization (Davidenko et al. 2012; Bracci and Op de Beeck 2016; Proklova et al. 2016). In addition, there are several segregated functional networks in human VTC that are specialized for processing different types of ecologically-relevant categories – such as faces and bodies (Haxby et al. 2000; Peelen and Downing 2007; Op de Beeck et al. 2008; Tsao and Livingstone 2008; Kanwisher 2010; Wandell et al. 2012; Grill-Spector et al. 2018). Indeed, previous findings in both humans and non-human primates have revealed causal relationships between neural responses in face-selective regions and the perception of faces (Afraz, Kiani, and Esteky 2006; Jonas et al. 2012; Moeller et al. 2017; Parvizi et al. 2012; Rangarajan et al. 2014), as well as between neural responses in body-selective regions and the perception of bodies (Downing and Peelen 2016). While these series of studies examined the relationship between neural responses and perception separately for each domain, it is presently unknown whether or how brain regions from these distinct cortical networks work together to achieve a behavioral goal such as perceptual categorization.

This gap in knowledge persists because previous research has largely focused on understanding how either (a) neural responses from one functional region contribute to perception (Hasson et al. 2001; Andrews et al. 2002; Moutoussis and Zeki 2002; Grill-Spector et al. 2004) or (b) neural responses are computationally transformed from regions that are positioned in early cortical stages of a network compared to later stages (Serre et al. 2007; Yamins et al. 2014). Building on this empirical foundation, a recent trend in neuroimaging studies has begun to unveil how information regarding different categories is combined. For example, several studies have shown that separate information about faces and bodies can be integrated within a functional region in VTC to generate new information that represents a whole person (Bernstein et al. 2014; Kaiser et al. 2014). While this approach sheds light on how information from these separate stimuli and both categories may be integrated in one functional region, it still remains untested (a) how regions selective for faces and bodies process stimuli that consist of visual features from both categories (Fig. 1A), and (b) if the combination of neural signals between regions explains behavioral categorization better than neural signals from either region by itself.

**Figure 1.**
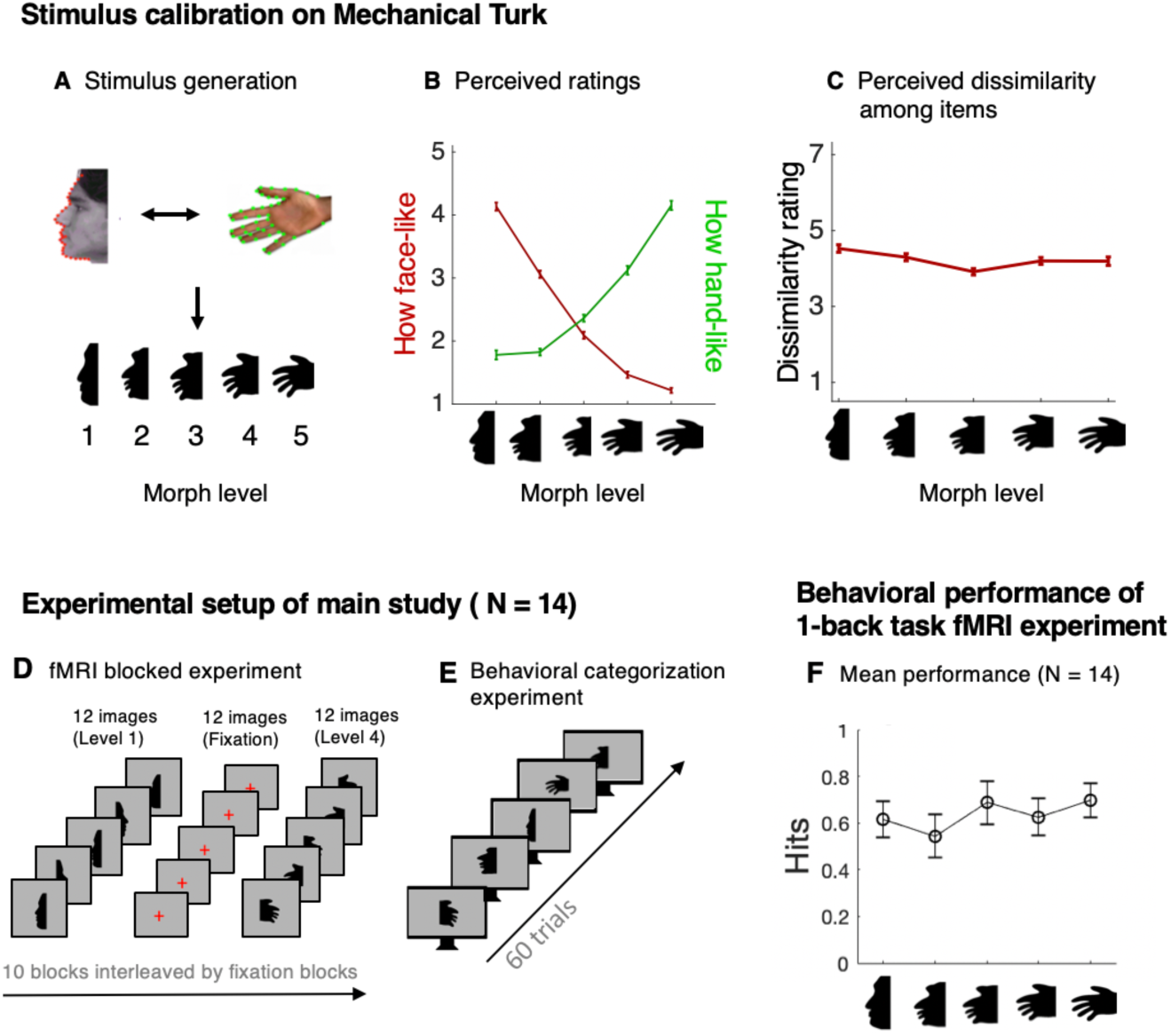
Stimulus calibration and experimental setup. **A.** Construction of face-hand silhouettes. *Top:* The contours of a profile face and open hand photograph are parameterized with 36 points in which points of maximum and minimum curvature are matched between faces and hands. *Bottom:* Five morph levels between faces and hands are created by interpolating the key point coordinates of parameterized faces and hands. The silhouettes are rendered by applying bi-cubic splines to the key points and shading the resulting figure in black. **B.** Morphed stimuli were calibrated on Mechanical Turk (MTurk) to produce intermediate stimuli (level 3) that were as face-like as they were hand-like (see Methods). *X-axis*: morph levels ranging from face (level-1, left) to hand (level-5, right). *Y-axis:* face-likeness ratings (red) and hand-likeness ratings (green) averaged across MTurk participants. *Errorbars:* Standard error across the 60 exemplars at each morph level. **C.** Matching perceptual variability. Each morph level consisted of multiple (12) exemplars. For each level, we calibrated the perceptual variability across those exemplars based on pairwise dissimilarity ratings to ensure equal variability of stimuli across all morph levels. *X-axis:* Morph level; *Y-axis*: Dissimilarity scores (higher scores indicate greater perceptual variability across the exemplars) averaged across MTurk participants. *Errorbars:* Standard error across 12 pairs of exemplars at each morph level. **D.** Each participant underwent a block-design fMRI experiment during which they viewed blocks containing images of carefully controlled silhouettes. 12 silhouettes from the same morph level were shown within each block. Subjects pressed a button when the same image appeared twice in a row (1-back task). **E.** Following the fMRI blocked experiment, each subject underwent a separate behavioral categorization experiment outside the scanner. Participants viewed silhouettes (N=60) and performed a forced-choice task to categorize each silhouette as a face or a hand. Presentation order was randomized across morph levels and participants. **F.** To control for attention during the fMRI block experiment, participants performed a 1-back task. For each level, we calculated the percentage of hits for each subject and averaged them for each morph level. Circles represent mean hit rate across N = 14 participants, while error bars represent standard error across subjects.

A rigorous way to begin to fill this gap in knowledge is with images that are generated on a known continuum of shape space in which (a) the ends of the continuum represent shapes from two different categories and (b) there is a clear categorical transition such that the center shape of the continuum is composed of equal parts of both categories (Fig. 1). Such a space enables a clear quantification of how neural and behavioral responses change when exposed to stimuli that cross category boundaries. Here, we generated such a shape space containing a morphed continuum between faces and hands using the silhouette methodology (Davidenko 2007; Davidenko et al. 2012) and tested if behavioral categorization of these stimuli is best predicted by either a) separate or b) combined neural responses from face- or body-selective regions in VTC. We reasoned that if the former model best predicts behavior it would support the idea that behavioral categorization of ambiguous stimuli is based on responses from a single region similar to prior results using natural, unambiguous stimuli (Andrews et al. 2002; Grill-Spector, Knouf, and Kanwisher 2004; Hasson et al. 2001; Moutoussis and Zeki 2002), whereas if the latter model best predicts behavior, it would suggest that behavioral categorization depends on relative neural responses across multiple regions despite the fact that these neural responses are from functionally distinct domain-specific networks.

## MATERIALS AND METHODS

### I. Generating and calibrating carefully-controlled face-hand morphs

We generated a novel set of parameterized silhouette stimuli that spanned a continuous morph space between faces and hands while controlling for many low-level image properties, including brightness, contrast, and total silhouette area. We generated 60 silhouette stimuli at each of 5 morph levels (ranging from fully face-like (level-1), ambiguously face- or hand-like (level-3), to fully hand-like (level-5)), which resulted in 300 stimuli. These stimuli were created by parameterizing the contours of a large set of photographs of profile faces and open hands using the same number of key points for images from both categories (Fig. 1A). Profile face photographs were obtained from the FERET database (Phillips et al. 1998, 1999) and were parameterized as outlined in Davidenko (2007), except that 36 rather than 18 key points were used to define each face contour. The increased number of key points compared to our prior work enabled a one-to-one mapping between the profile face photographs with the more detailed hand contours. Open hand photographs were obtained from 12 volunteers who each provided between 10 and 20 open hand poses, for a total of 148 unique hand images. As with the face images, 36 key points were defined along the contours of each hand, such that points of maximum and minimum curvature matched between faces and hands (e.g. point 7 corresponds to the tip of the brow in faces and the tip of the thumb in hands; see Fig. 1A).

#### Stimulus Calibration

The silhouette stimuli were calibrated in an iterative process by (1) obtaining perceptual ratings from Mechanical Turk (mTurk) workers (https://www.mturk.com/), (2) adjusting the parameterized stimuli, and (3) repeating the process. This calibration procedure had two goals: first, to obtain intermediate (level-3) stimuli that appeared equally face- and hand-like, and second, to generate multiple exemplars at each morph level that were matched for perceptual variability across the five morph levels. To accomplish the first goal, mTurk workers rated between 1 and 10 stimuli randomly selected from among the 300 exemplars (60 at each morph level) on either how face-like (1 to 5 scale) or how hand-like (1 to 5 scale) they appeared, until each exemplar had been rated by at least 5 workers. Following this first round, intermediate stimuli were found to be too hand-like, so these were adjusted in the face-hand morphing trajectory toward faces. The second round produced improved results, and the third round produced stimuli that were well-balanced across the five morph levels (see Fig. 1B). To control for perceptual variability, a similar iteration of mTurk studies was conducted in which workers rated between 1 and 10 pairs of stimuli from within each of the 5 morph levels, according to a 7-point dissimilarity metric (1=identical and 7=maximally dissimilar). After the first round, level-1 (face) stimuli were found to be more perceptually variable than the other 4 levels, and so their variability was reduced for the second round by morphing each face stimulus 40% toward the average face. The second round of mTurk ratings produced well-matched perceptual variability across the five morph levels (Fig. 1C).

### II. Testing categorization of face-hand morphs within and outside the scanner

#### Participants

14 participants (8 females, ages 22 - 44) were scanned at the Center for Neurobiological Imaging (CNI) at Stanford University using a 3 T GE Discovery MR750 scanner. Each subject underwent (1) an anatomical scan, (2) a functional localizer experiment, (3) a face/hand morphing experiment within the scanner (Fig. 1D), and afterwards, (4) a behavioral experiment outside the scanner that measured their categorization of face/hand morphs as either faces or hands (Fig. 1E). All participants gave their written informed consent. Procedures were approved by the Stanford Internal Review Board on human participant research.

##### 1. Anatomical imaging and cortical surface reconstruction

###### Data acquisition

A standard high-resolution anatomical volume of the whole brain was acquired using a T1-weighted BRAVO pulse sequence (TR=450 ms, flip angle=12°, 1 NEX, FOV=240 mm, resolution: 1.0mm isotropic).

###### Cortical surface reconstruction

All anatomical volumes were aligned to the AC-PC plane. Using automated (FreeSurfer: http://surfer.nmr.mgh.harvard.edu) and manual segmentation tools (ITK-SNAP: http://www.itksnap.org/pmwiki/pmwiki.php), each anatomical volume was segmented to separate gray from white matter, and the resulting boundary was used to reconstruct the cortical surface for each subject (Wandell et al. 2000).

##### 2. Functional Imaging

###### Data acquisition

Using a custom-built phase-array, 32-channel head coil, we acquired 34 oblique coronal slices covering occipito-temporal cortex (resolution: 2.4 mm x 2.4 mm x 2.4 mm; one-shot T2*-sensitive gradient echo acquisition sequence: FOV = 192 mm, TE = 30 ms, TR = 2000 ms, flip angle = 77°).

###### Localizer Experiment

All subjects participated in 2 runs of an fMRI functional localizer experiment (150 volumes each). Each run consisted of 12 s blocks, which contained different images from the same category presented at a rate of 1 Hz. Images within each block were from 1 of 7 different categories: faces (frontal / profile view), body parts (hands / lower limbs), objects (cars / tools), and scrambled versions of these images. Across the two experimental runs, there were 8 repetitions of each block type. Participants maintained fixation and responded by button press when an image repeated (1-back task).

###### Morphing experiment

Using the carefully calibrated face and hand morph stimuli, all subjects participated in 4 runs of a block design experiment during which subjects viewed silhouettes alternating with a gray screen. A fixation cross was on the screen at all times (Fig. 1D). Each run consisted of 10, 12-second blocks and each consisted of 12 silhouettes displayed at a frequency of 1 Hertz. Blocks of silhouettes (a) always contained images from the same morphing level and (b) appeared in a pseudo-random order, in which there was a difference of two morphing levels (on average) between consecutive blocks (e.g. level 3 followed by level 1, or level 2 followed by level 4). In order to assure that any measurable effects in high-level visual regions were not driven by consistent placement of silhouette edges in a particular location in the visual field (Davidenko et al. 2012), (1) images were placed centrally in one of 8 randomly-chosen locations around the fixation point (max visual degree: 3), and (2) silhouettes faced either left or right. During the silhouette blocks, participants maintained fixation and responded by button press when an image repeated (1-back task), which happened 0-2 times per stimulus block. In order to avoid fMRI adaptation effects, the same silhouette was never repeated other than for a 1-back task trial. To keep task and cognitive processes as similar as possible between stimulus and rest blocks, fixation crosses also changed with a 1 Hertz frequency displaying different lengths of lines during the rest blocks, and participants performed the 1-back task when the same fixation cross appeared twice in a row. Analyses of 1-back performance during task blocks revealed that participants’ performance was similar across blocks from all morph levels (Fig, 1F). For each subject and each morph level, we computed the hit rate by dividing the total number of hits by the total number of 1-back occurrences for a given morph level. False alarm rates were 2 % or lower for each morph level and were therefore not taken into account. A non-parametric repeated measures analysis of variance (Friedman’s test) of correct 1-back responses revealed marginally significant differences across morph levels (Chi-squ(4,52) = 8.96, *p* = 0.06).

##### 3. Behavioral experiment of face/hand morph categorization

To ensure that subjects were blind to the behavioral task during the fMRI experiment, each of the 14 subjects also participated in a behavioral testing experiment outside the scanner *a few weeks after* the fMRI experiment. The experiment was self-paced and consisted of 60 trials using the same silhouettes used during the fMRI experiment (12 images from each of the 5 morph levels, presented in random order). In each trial, a silhouette appeared at the center of the screen (Fig. 1D). Subjects then performed a forced-choice task, deciding whether the silhouette was a face (rating: 1) or a hand (rating: 0). The stimulus remained on the screen until the subject chose one of the two options and no additional instructions were given. For each of the subjects, we recorded the mean response for each morphing level, resulting in 5 values between 0 (all hand) and 1 (all face) for each subject.

### III. Data analyses

fMRI data were analyzed using mrVista (http://github.com/vistalab) and custom software written in and implemented by MATLAB (www.mathworks.com). Within-subject analyses were performed for each of the 14 subjects.

#### 1. Timeseries preprocessing

Functional data were motion corrected using an affine transformation (Nestares and Heeger 2000). Time series data were processed with a temporal high-pass filter (1/20 Hz cutoff) and then converted to percentage signal change by dividing the time series of each voxel by its mean intensity. We estimated the blood oxygen level dependent (BOLD) response amplitudes for each condition using a general linear model (GLM) applied to the time series of each voxel. Predictors of the GLM were the experimental conditions convolved with the hemodynamic impulse response function used in SPM (http://www.fil.ion.ucl.ac.uk/spm/). Data were not spatially smoothed.

#### 2. Functional regions of interest (ROIs)

Our analyses focused on functional regions of interest (ROIs) that are selective for faces and limbs within ventral temporal cortex (VTC) as VTC is critical for categorization (Grill-Spector and Weiner 2014) and participants performed a categorization task during the behavioral experiment.

##### Face-selective ROIs

Face-selective voxels were identified as those voxels in VTC that illustrated higher BOLD responses to grayscale photographs of faces (frontal and profile view) compared to body parts (hands and lower limbs) and objects (cars and tools) in the functional localizer experiment (Fig. 2A). Profile views of faces were originally included in the localizer to test if different functional regions resulted from different statistical contrasts: 1) profile face images > objects and bodies and 2) frontal faces images > objects and bodies. Results showed that both contrasts produced the three typical face-selective regions in VTC (IOG-faces, pFus-faces, mFus-faces). Thus, blocks of profile and frontal face images were combined in the statistical contrast used for localization. As in our prior publications, two VTC ROIs were defined within each subject’s native brain anatomy using a common threshold (*t* > 3, voxel-level) and were positioned lateral to the mid-fusiform sulcus (MFS; Weiner et al., 2014). mFus-faces is adjacent to the anterior tip of the MFS, while pFus-faces is located 1-1.5cm more posteriorly on the lateral fusiform gyrus extending into the posterior occipitotemporal sulcus (OTS). In some studies mFus-faces is also referred to as FFA-2, while pFus-faces is also referred to as FFA-1, in which the FFA stands for Fusiform Face Area (Kanwisher et al. 1997). We list which face-selective regions in VTC were identifiable in each subject in Supplemental Table 1, as well as the stereotaxic coordinates (from Weiner and Grill-Spector, 2013) of each region in Supplemental Table 2.

**Figure 2.**
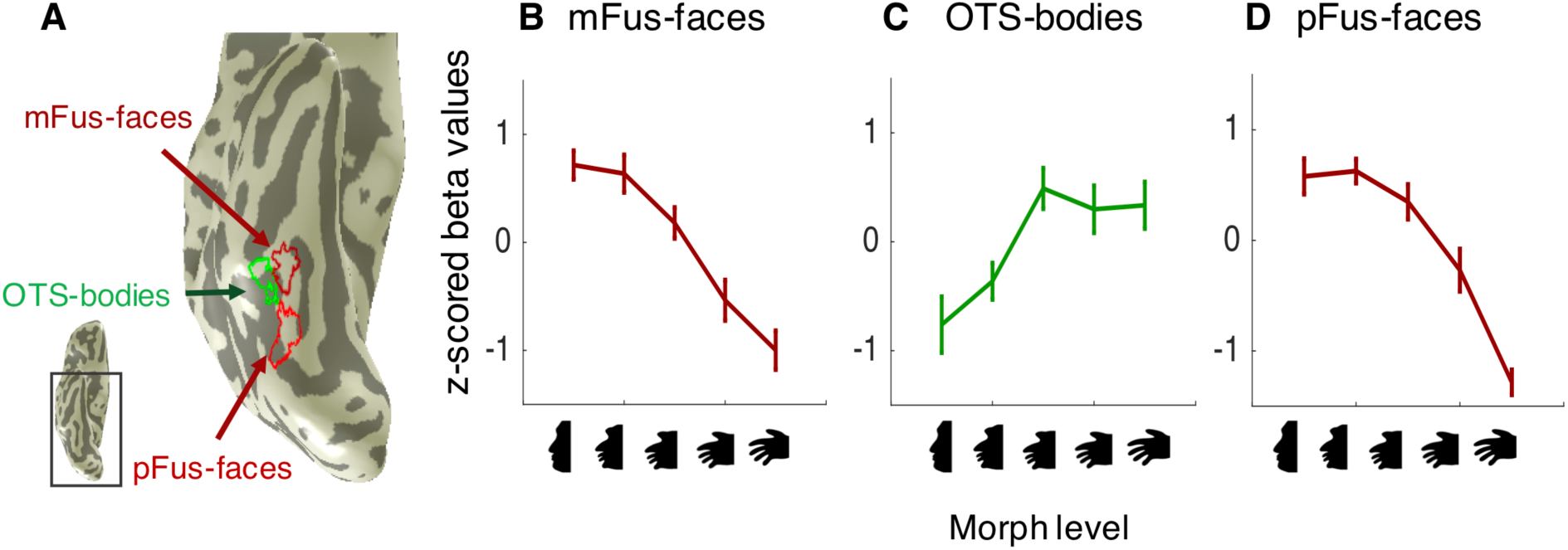
Face- and body-selective regions in ventral temporal cortex (VTC) display differential neural tuning to face-hand morphs. **A.** Zoomed image showing the inflated cortical surface in ventral view of an example participant illustrating three regions of interest (ROIs). From posterior to anterior: pFus-faces (red), OTS-bodies (green), and mFus-faces (dark red). *Dark gray:* sulci. *Light gray*: gyri*. Inset:* Ventral view of the inflated cortical surface showing the location of the zoomed region. **B-D.** The mean neural tuning across participants for each fROI of the right hemisphere. Normalized responses were derived by first computing the beta coefficients to each one of the 5 morph levels of face-hand silhouettes and then transforming the beta coefficients into z-scores. *X-axis*: face-hand silhouette morph level. *Y-axis:* Z-scored beta values. *Errorbars:* Standard error across participants. See **Supplemental Table 1** for the number of participants per ROI (N=11-14). See **Supplemental Fig. 2** for left hemisphere data.

##### Body-selective ROI

Body-selective voxels were identified as those voxels in VTC that illustrated higher BOLD responses to grayscale photographs of limbs (hands and lower limbs) compared to faces (frontal and profile view) and objects (cars and tools) in the functional localizer experiment. As in our prior publications, one ROI was defined within each subject’s native brain anatomy using a common threshold (*t* > 3, voxel-level) and was located within the OTS, positioned between mFus-faces and pFus-faces. We refer to this region as OTS-bodies, but it is also referred to as the Fusiform Body Area (Schwarzlose et al. 2005; Peelen and Downing 2006). We list which body-selective regions were identifiable in each subject in Supplemental Table 1, as well as the stereotaxic coordinates (from Weiner and Grill-Spector, 2013) of each region in Supplemental Table 2.

#### 3. Mean tuning analyses

##### Neural tuning

Within each individual subject, a GLM was conducted on the mean fMRI timeseries of each ROI (mFus-faces, pFus-faces, OTS-bodies), separately for the left and right hemisphere. Response amplitudes (betas) and residual variance of each ROI were estimated from the GLM, which yielded 5 beta values per ROI (one for each morphing level; Fig. 2). Beta values were z-scored (mean: 0, SD: 1) to allow for comparison across subjects and ROIs. For each ROI, main effects of morph level were separately evaluated using a 1-way repeated measures analysis of variance (ANOVA) with morph level as a factor. The interaction between morph level, ROI, and hemisphere was also evaluated using a repeated measures ANOVA using morph level, ROI, and hemisphere as factors. We refer to the relationship among these five beta values as *neural tuning* to these face-hand stimuli.

##### Behavioral Tuning

For each subject, we measured the average categorization responses of each morph level across the 12 trials of that level during a behavioral experiment outside the scanner. This results in one behavioral response for each of the 5 morph levels, indicating the proportion of face responses for the given morphing level. We refer to this function as *behavioral tuning* to these face-hand stimuli (Fig. 3).

**Figure 3.**
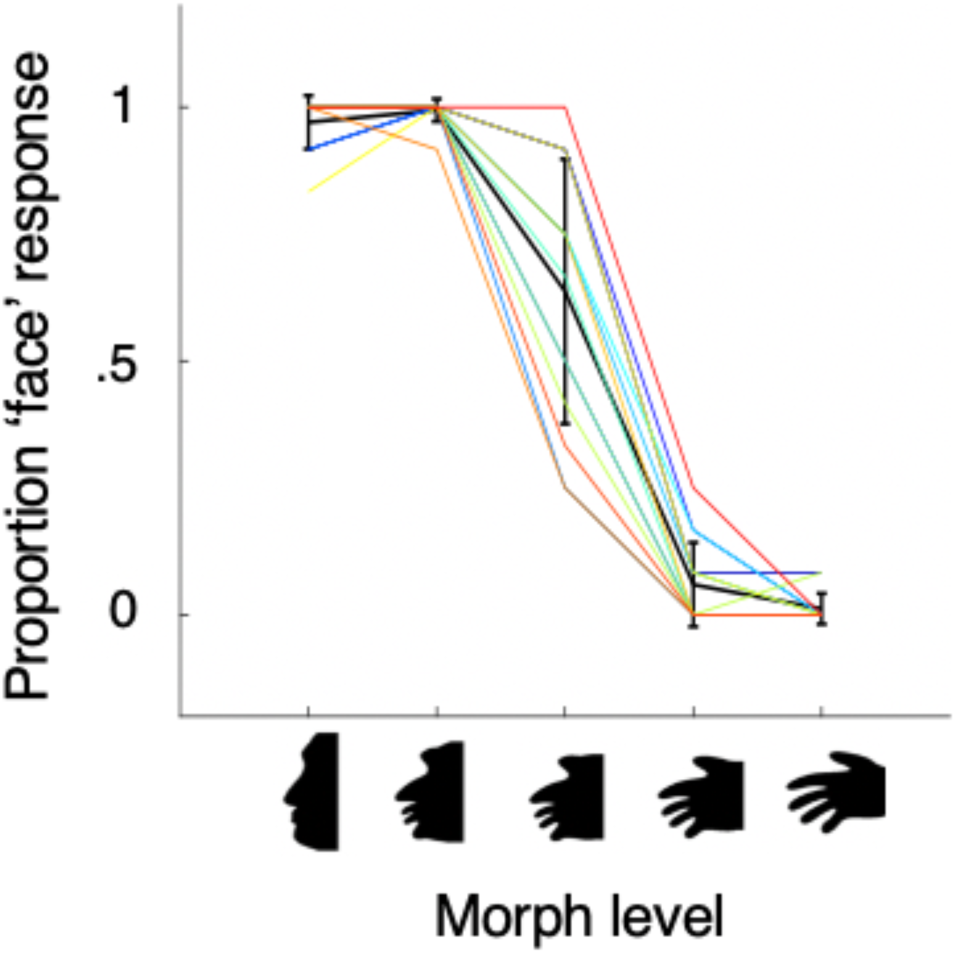
Behaviorally, participants effectively categorize the face-hand morphs. Behavioral tuning for each subject (colored lines) as well as the mean tuning across subjects (black line). Behavioral responses at each morph level indicate the participant’s average performance across 12 trials. Proportion ‘face’ responses decline with morph level showing a categorical transition. The highest variability in behavioral responses across participants occurred for the center morph level. *X-axis:* morph level of silhouette (1 = face, 5 = hand). *Y-axis:* proportion face response. *Errorbars:* Standard deviation across the 14 participants.

#### 4. Relating neural tuning to behavioral tuning

In order to quantify the relationship between neural tuning and behavioral tuning, we used a linear regression model that relates each subject’s behavioral tuning to their neural tuning. Each ROI’s tuning (z-scored beta values) to the five morph levels was used as a predictor for a linear regression model (the independent variable) relating it to the behavioral data (the dependent variable, Fig. 4A). This analysis was performed on a subject-by-subject basis and separately for neural data from each hemisphere. Goodness-of-fit of the model was evaluated by computing the explained variance (R^2^). R^2^ was calculated as the square of the correlation between the model’s estimation (1 x 5 vector) and the behavioral data (1 x 5 vector). Finally, simulations of such a 5-point by 5-point model provide a null distribution that guided the evaluation of our modeling results (Supplemental Fig. 1). We used paired permutation testing (10,000 iterations) to assess whether the explained variances (R^2^) differed significantly between model pairs. For example, comparing the OTS-bodies model and mFus-faces models. Specifically, for each iteration and each subject, we randomly permuted the two model labels. Then, the null distribution is defined as the distribution of R^2^ differences (between the two models) using these permuted labels. This allows us to assess what differences in R^2^ values we could expect by chance. Consequently, we are able to compare our measured R^2^ differences across models to this null distribution. Permutation tests were used because we did not want to make any assumptions about the distribution of R^2^ difference values. Results were corrected for multiple comparisons using the Bonferroni method (dividing the p-value by the number of comparisons minus one). For each permutation test, we only used those subjects that had the ROIs included in both models (Supplemental Table 1). Permutation tests for all analyses in this article are performed with the same methodology.

**Figure 4.**
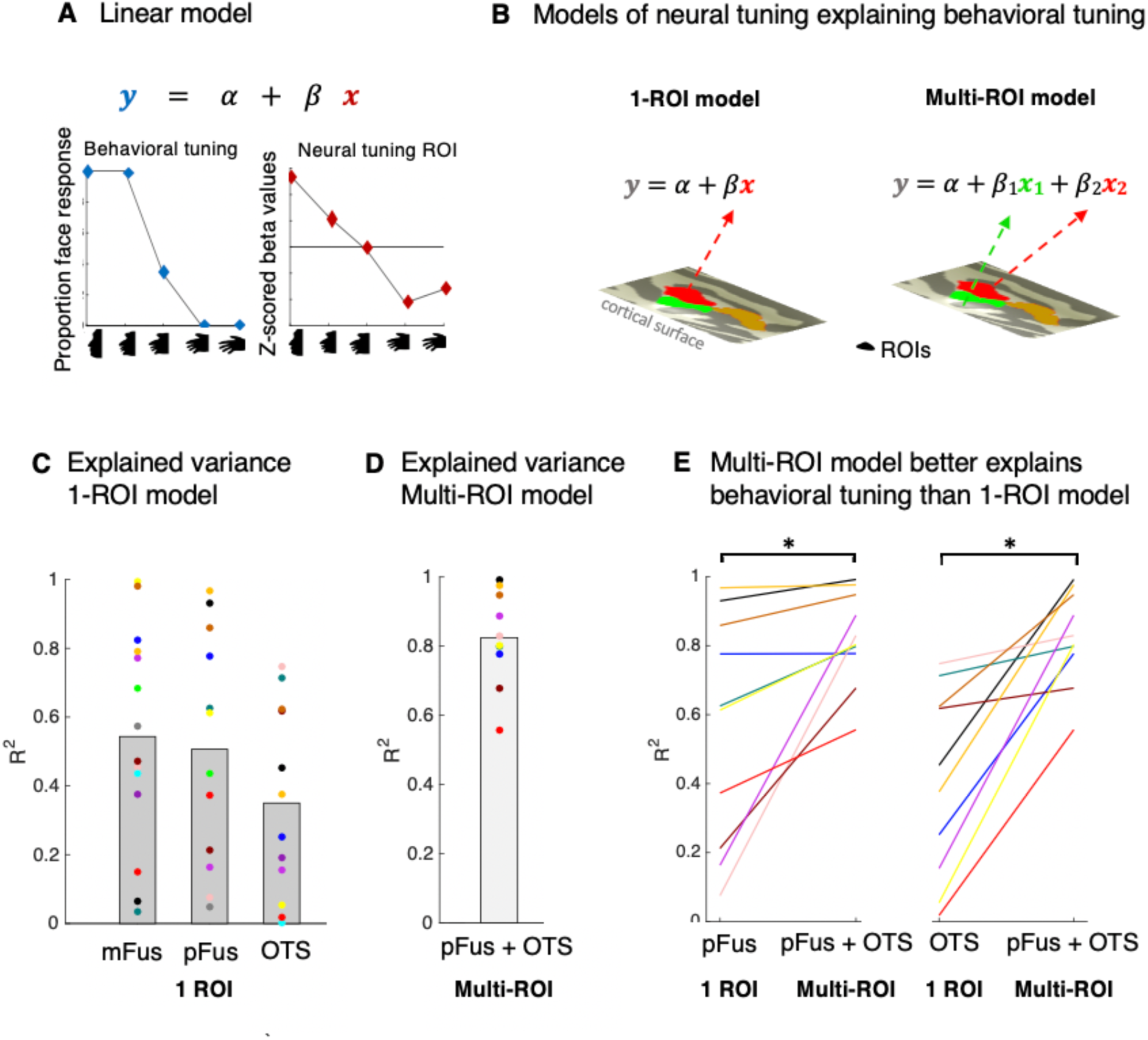
A stepwise linear regression model reveals that behavioral tuning to face-hand morphs can be predicted by a linear combination of the neural tuning from pFus-faces and OTS-bodies. **A.** *Top:* A linear regression model was used to relate behavioral and neural turning. *α* = model intercept, *β* = beta coefficient for predictor, x =neural predictor, y = behavioral tuning. *Left:* Behavioral tuning. *Right:* Neural tuning. **B.** We tested two different types of linear models: (1) a 1-ROI model, which used neural tuning from one ROI as a predictor for behavioral tuning, and (2) a multi-ROI model (MRM), which used neural tuning from multiple ROIs as separate predictors based on results from a stepwise linear regression. **C.** Explained variance (R^2^) from the 1-ROI model in the right hemisphere. Neural tuning from each ROI was a significant predictor of behavioral tuning. *X-axis:* ROIs used as predictors for the linear regression model. *Y-axis:* Explained variance of the behavioral data by three 1-ROI models. Bars indicate the mean across participants, and each dot represents data from one participant. The color of each participant is matched across models and data in Fig. 3. **D.** Explained variance of the behavioral data by the MRM based on neural tuning from right pFus-faces and right OTS-bodies. Conventions same as C. **E.** Improvement in explained variance by the MRM compared to the 1-ROI model. Each line is a participant. Line colors correspond to (C). *Asterisk*: significant improvement, *p*<.05, permutation testing.

##### Linear regression model for individual ROI data (1-ROI model)

We implemented 6, 1-ROI linear regression models predicting the behavioral tuning curve from each of the ROIs (mFus-faces, pFus-faces, OTS-bodies; separately for LH and RH, Fig. 4B).

##### Linear regression model using data from multiple ROIs (Multi-ROI model, MRM)

We ran one stepwise linear regression model using all subjects and the three ROIs (mFus-faces, pFus-faces, OTS-bodies; separately for each hemisphere and concatenated across subjects) to establish which of the ROIs, or which combination of ROIs, best explained behavioral tuning. After identifying which regions were significant predictors, we applied a linear regression model that used data from pFus-faces and OTS-bodies as separate predictors for behavioral tuning as these were the ROIs that were identified as significant predictors during the stepwise linear regression (Fig. 4B). As in the 1-ROI model, this was done individually for each subject. We note that we could identify both pFus-faces and OTS-bodies in 10/14 subjects. Thus, further analyses with right hemisphere pFus-faces and OTS-bodies were performed using these 10 subjects.

To test the improvement by the MRM compared to the 1-ROI model, we used the model selection criterion AIC (Akaike’s Information Criterion) to compare model performances, in which lower values indicate a better model fit. In addition, we used paired permutation testing (10,000 iterations) to examine whether the Multi-ROI model outperformed the 1-ROI models. Finally, we compared if there were any hemispheric differences in model performance using paired permutation testing (using multiple comparison correction as described above).

##### Linear regression Multi-ROI control models

We considered two Multi-ROI control models. First, to test if any 2-ROI model can predict behavioral tuning to face/hand morphs, we ran a control model using 2 ROIs that are cortically proximal to face- and limb-selective regions, but are not selective for either faces or limbs. These ROIs were a retinotopically defined visual area (VO-2 from Wang et al., 2014) and a place-selective region (CoS-places from Weiner et al., 2018). Both regions were from published atlases and were projected to individual subject space using FreeSurfer’s cortex-based alignment (CBA) as in our previous work (Rosenke et al. 2017; Weiner et al. 2018). Analyses were performed in the same fashion as the Multi-ROI model described above.

Second, to test if the 2-ROI model was specific to neighboring face- and body-selective regions in VTC or generalized to lateral occipito-temporal cortex (LOTC), in which face- and body-selective regions also neighbor one another (Orlov et al. 2010; Weiner and Grill-Spector 2011, 2013), we defined face- and body-selective ROIs in LOTC. We defined a face-selective region on the inferior occipital gyrus (IOG-faces), which is also known as the occipital face area. We also defined three body-selective regions on the middle temporal gyrus (MTG-bodies), the inferotemporal gyrus (ITG-bodies), and the lateral occipital sulcus (LOS-bodies), which are often described as one extrastriate body area (Downing et al., 2001), but are anatomically and functionally distinct (Weiner and Grill-Spector, 2011, 2013). To mirror our analyses for VTC category-selective regions, we ran a stepwise linear regression using data from face- and body-selective regions in LOTC (Supplemental Table 1 for the frequency of face-selective and body-selective regions identifiable in each subject within LOTC). Further analyses and comparisons to the VTC Multi-ROI model are done as described in the previous sections.

#### 5. Cross-Validated MRM model across participants

We tested if a model built from the neural responses in one group of participants could predict behavioral tuning in a new, independent participant. For these analyses, we used neural tuning from pFus-faces and OTS-bodies because our previous linear regression analyses indicated that neural tuning from these regions were the best predictors of behavioral tuning.

We first estimated regression weights using a linear regression analyses based on data from N-1 subjects and then predicted responses of the left-out subjects (Arlot and Celisse 2010; Breiman and Spector 1990). This analysis was done using 10 subjects in which we could define both of the ROIs (see Supplemental Table 1). To perform the analysis, we concatenated the z-scored neural responses to the 5 morph levels in each ROI across subjects, which resulted in a 1×45 (e.g. 9 subjects x 5 morph levels) vector per ROI. Likewise for behavioral responses, we concatenated the behavioral responses to the 5 morph levels across subjects (1×45 vector). The two neural vectors served as the predictors for the behavioral vector. The regression analysis derived the weights (model coefficients) for each of the two neural predictors. To predict responses of the left-out subjects, we multiplied the model coefficients by the measured neural responses of the left-out-subject to generate the predicted behavioral tuning for the left-out-subject. We then calculated the goodness-of-fit between the predicted and actual behavioral tuning of the left-out-subject using R^2^ as described previously. This procedure was cross-validated using all 10 subjects, resulting in a 10-fold cross-validation. This cross-validated Multi-ROI model was compared to the two cross-validated Control Models, which used (1) neural data from VO-2 and CoS-places as predictors and (2) neural data from IOG-faces and ITG-bodies, which were selected using the MRM stepwise linear regression model implemented with LOTC ROIs. The models were compared using paired permutation testing as well as multiple comparison correction as described in previous sections.

## RESULTS

We first generated 60 face-hand morphs using the silhouette methodology (Davidenko 2007). Compared to natural images, the silhouette methodology holds the advantage of generating a stimulus space that is a) controlled for low level visual features, b) changes only one visual attribute of the stimulus (shape), and c) drives neural responses in category-selective regions in VTC (Davidenko et al. 2012). This approach generated a large set of carefully controlled images consisting of continuous morphs between faces and hands (Fig. 1A; Materials and Methods).

We calibrated the stimuli on Mechanical Turk (www.mturk.com). Calibration results indicated that (a) the morph continuum was centered, which means that the middle morph level was ambiguous and equally likely to be perceived as either a face or a hand by individual observers (Fig. 1B), and (b) the perceptual variability across exemplars within each morph level was similar (e.g., level 1 exemplars were perceived as being as variable as level 4 exemplars, Fig. 1C). The latter is important as the lack of variability of stimuli within a morph level may generate fMRI-adaptation (Davidenko et al. 2012; Grill-Spector and Malach 2001). Thus, our calibration ensured that the variability of stimuli is matched across morph levels.

Using these stimuli, we conducted two types of experiments in 14 independent adults (8 females, ages 22 - 44): (1) A functional magnetic resonance imaging (fMRI) block-design experiment (using a 1-back task as an attentional control) in which we measured mean neural responses to face-hand morphs along the morphing continuum in face- and body-selective regions in VTC (Fig. 1D). We refer to neural responses across morph levels as neural tuning. (2) A behavioral categorization experiment outside the scanner during which we measured the proportion of face-hand morphs in each level that are perceived as faces or hands, which we refer to as behavioral tuning (Fig. 1D).

### Face- and body-selective regions in ventral temporal cortex (VTC) are tuned to a continuum of face- and hand-like silhouettes

We first asked how functional regions of interest (ROIs) identified in VTC with natural images respond to ambiguous face-like and hand-like silhouettes, respectively. To do so, we identified face- and body-selective regions within VTC: (1) pFus-faces/FFA-1, which is a face-selective region located in the posterior-lateral portion of the fusiform gyrus (FG; Kanwisher et al. 1997; Weiner and Grill-Spector 2010; Weiner et al. 2014)), (2) OTS-bodies/FBA, which is a body-selective region located within the occipitotemporal sulcus (OTS; Peelen and Downing 2005; Schwarzlose et al. 2005; Weiner and Grill-Spector 2010)), and (3) mFus-faces/FFA-2, which is a face-selective region located within the FG extending laterally from the anterior tip of the mid-fusiform sulcus (Kanwisher et al. 1997; Weiner and Grill-Spector 2010; Weiner et al. 2014; Fig. 2). From these ROIs, we then extracted neural responses (z-scored beta values from general linear model fits; Materials and Methods) to the morphed silhouettes that were acquired during a second, independent experiment (Fig. 1D).

Qualitatively, both pFus- and mFus-faces showed differential responses, or neural tuning, to face-hand morphs. pFus- and mFus-faces showed the highest neural responses to face-like silhouettes and the lowest responses to hand-like silhouettes. Interestingly, responses were a) similar across the two morph levels that were more face-like and b) only declined past the center morph level. Comparatively, OTS-bodies showed a different tuning profile in which the highest neural responses were to hand-like silhouettes and the lowest responses were to face-like silhouettes. Additionally, responses in OTS-bodies were a) similar across morph levels that were more hand-like, as well as the center morph level, and b) only declined in response for the two morph levels that were more face-like (Fig. 2B,D and Supplemental Fig. 2A,C). These observations were statistically supported by a 3-way repeated measures ANOVA with ROI, morph level, and hemisphere as factors (morph level x ROI interaction (F (8,305)=16.45, p <0.001; no significant difference between hemispheres (F (1,168) = 5.6458e-29, p = 1.0). Follow-up 1-way ANOVAs within each ROI (Bonferroni corrected) further supported an effect of morph-level (mFus-faces: F (4,120) = 5.08, p<0.005; pFus-faces: F (4,110) = 4.52, p<0.01; OTS-bodies: F (4,80) = 5.93, p<0.001). Together, these analyses reveal that a) face- and body-selective regions in VTC display neural tuning to ambiguous face-hand silhouettes and b) this tuning is not linear with the morphing continuum.

### Categorization behavior of face-hand morphs is best explained by neural tuning from multiple category-specific regions from different domains

Since face- and body-selective regions in VTC display selective tuning to face-hand silhouettes, we asked: *What is the best neural predictor for perceptual categorization of these ambiguous face-hand stimuli?* To answer this question, we first used the same face-hand silhouettes that participants previously viewed inside the scanner to measure how our participants perceptually categorized the same stimuli outside the scanner. In the behavioral experiment, participants viewed 60 silhouettes randomly drawn from the 5 morph levels (12 exemplars from each morphing level) and were instructed to classify each image as either a face or a hand, in a self-paced manner (Fig. 1E). We refer to behavioral responses relating these forced-choice behavioral responses to the morph levels as *behavioral tuning* (see Methods).

Behavioral results showed that participants perceptually differentiated face from hand silhouettes. That is, participants classified the two more face-like morph-levels as faces in the majority of trials (Fig. 3, proportion face responses (mean±standard deviation (SD)): .97±.05 and .99±.02, respectively), while they classified the two more hand-like morph levels as hands (proportion face response: .06±.08 and .01±.03, respectively). Categorization of the middle morph level was more variable across participants. On average, images in the middle, or intermediate, morph level 3 were classified slightly more as faces (proportion face response: .64±.26).

We next tested if and how behavioral judgements are linked to neural responses in face- and body-selective regions. To do so, we conducted a linear regression analysis relating the behavioral categorization data to neural responses in face- and body-selective ROIs. In this analysis, the behavioral tuning of each subject is the dependent variable and the neural tuning of that subject is the independent variable (Fig. 4A). We considered two models: (1) a linear regression model relating behavioral categorization to neural responses from a single face- or body-selective region (Fig. 4B**, 1-ROI model**) and (2) a linear regression model relating behavioral categorization to neural responses from multiple regions (Fig. 4B**, Multi-ROI model**). For the latter, we implemented one version of the model that contained neural responses from two regions selected by a stepwise linear regression model. In addition, we implemented two control models using neural responses from a) two regions that were neither face- nor body-selective and b) two regions that were selective for faces and bodies not located in VTC, and instead, in LOTC.

Results of the 1-ROI model showed that on average, behavioral categorization performance from individual participants was linearly related to neural responses from either face- or body-selective regions. That is, neural responses from either face (pFus-faces, mFus-faces) or body (OTS-bodies) ROIs explained, on average, a substantial amount of the variance of behavioral responses (R^2^± standard error across subjects (SE), left hemisphere (LH) and right hemisphere (RH), respectively: mFus-faces = .43±.10 and .54±.09; pFus-faces = 47±.09 and .51±.09; OTS-bodies = .41±.06 and .35±.07; RH: Fig. 4C; LH: Supplemental Fig. 3A). However, there was extensive variability in model fits across participants whereby in some participants the variance explained was close to 1, but in others, it was close to 0 (R^2^ min-max: 0 - .99; Fig. 4C). This between-subject variability in model fits suggests that the 1-ROI model is not a parsimonious model of behavioral categorization.

We next implemented a multiple ROI model in which we first used a stepwise linear regression model to test which combination of ROIs best explains behavioral tuning. The stepwise regression was applied to data from all subjects and neural responses from all three ROIs (Fig. 4B). Interestingly, this model revealed that neural tuning from two ROIs: pFus-faces (LH and RH, *p*s<0.001) and OTS-bodies (LH and RH, *p*s<0.001) significantly explained behavioral categorization. Adding mFus-faces to the model did not significantly explain additional variance (LH and RH: *p*s = 0.18). Consequently, the remaining analyses only consider a model of behavioral categorization using pFus-faces and OTS-bodies as predictors, which we refer to as the Multi-ROI model (MRM). As only 10 out of 14 subjects had both pFus-faces and OTS-bodies across hemispheres, subsequent analyses are constrained to those 10 subjects (Supplemental Table 1).

Examining the MRM fit for each individual participant showed that the MRM model explained 0.71±.03 and 0.83±.04 (R^2^±SE) of the variance of behavioral categorization from neural responses in the left and right hemispheres, respectively (Fig. 4D, Supplemental Fig. 3B). Comparing the variance explained by the MRM to the 1-ROI model revealed higher variance explained by the former than the latter in each participant (Fig. 4E). As a two vs. one parameter model comparison results in different degrees of freedom, we tested the statistical significance of these data in two ways: (1) Akaike’s Information Criterion (AIC) and (2) paired permutation tests between each of the 1-ROI models and the MRM. Mean AIC (mean across subjects ± SE for LH, RH) was as follows: pFus-faces = 5.40±1.24, 4.55±1.46; OTS-bodies = 6.96±0.57, 7.27±0.63; pFus-faces + OTS-bodies = 5.19±0.66, 0.88±1.69. As lower AIC values indicate better model fits, the data suggest that a linear combination of neural responses of the right pFus+OTS best explains behavioral categorization of face-hand silhouettes. Additionally, paired permutation tests showed that significantly more variance of behavioral categorization was explained by the MRM (all ps<0.05, Bonferroni corrected, Fig. 4E) than each of the 1-ROI models. Differences in model predictions across hemispheres are not significant (paired permutation tests between left and right hemisphere model: *p*s>0.05).

While these results are clear, it may just be the case that neural signals from multiple ROIs – irrespective of their selectivity – better explain behavioral categorization compared to responses from a single ROI. To test this possibility, we implemented a control MRM that related behavioral categorization to neural tuning from two other functional regions not selective for faces or bodies. For this model, predictors were neural responses from ROIs that either did not illustrate a specific categorical preference (retinotopic area, VO-2 (Fan et al. 2014)), or were not selective to either faces or hands (Parahippocampal Place Area (PPA)/CoS-places (Aguirre et al. 1998; Epstein and Kanwisher 1998; Weiner et al. 2018)). This model had a significantly (permutation test, *p* < .05) lower average explained variance (R^2^±SE: .52±.06 (LH) and .58±.08 (RH), Supplemental Fig.3C) and a higher AIC criterion (7.64±0.79 (LH) and 5.48±1.55 (RH)) compared to the MRM model. Interestingly, this control MRM was also not significantly different from any of the 1-ROI models (adjusted Bonferroni correction within each hemisphere, permutation testing. all p > 0.10).

We implemented an additional control model to test if the face- and body-selective MRM was specific to neighboring face- and body-selective regions in VTC, or generalized to lateral occipito-temporal LOTC, in which face- and body-selective regions also neighbor one another (Orlov, Makin, and Zohary 2010; Weiner and Grill-Spector 2013). Similar to how we established the ROI combination for the MRM model in VTC, we ran a stepwise linear regression model using face-selective (IOG-faces) and body-selective ROIs (LOS-bodies, MTG-bodies, and ITG-bodies) in LOTC. Interestingly, the stepwise linear regression model showed that the results generalized to LOTC in which a specific combination of neural responses explained more variance than other combinations: cortically adjacent ROIs, ITG-bodies (LH and RH: p< 0.0001) and IOG-faces (LH: p < 0.005, RH: p < 0.0001), explained a significant amount of variance of behavioral tuning, while other ROIs (LOS-bodies, MTG-bodies, *p*’s > 0.5) did not. Interestingly, explained variance (R^2^±SE: 0.80±0.05 (LH) and 0.82±0.05 (RH)) of the LOTC MRM was comparable to, and did not significantly differ from, the VTC MRM (permutation testing LH: p = 0.23, RH: p = 0.45, Supplemental Fig. 3). As the MRM model generalized to face-and body-selective regions in LOTC, the LOTC MRM will be included in further analyses.

A closer look at the VTC MRM and LOTC MRM revealed that the model-estimated behavioral tuning closely matched the measured behavioral tuning of a given participant (Fig. 5, compare to 1-ROI model estimations in Supplemental Fig. 4). Furthermore, the model consistently assigned opposing weights to pFus-faces (LH: β_pFus_ = 0.22±.06, RH: β_pFus_ = 0.39±.06) and OTS-bodies (LH: β_OTS_ = -0.22±.05, RH: β_OTS_ =-0.22±.07, Fig. 5**, right column**), as well as IOG-faces (LH: β_IOG_ = 0.39±.08, RH: β_IOG_ = 0.32±.09) and ITG-limbs (LH: β_ITG_ = -0.47±.11, RH: β_ITG_ = -0.28±.08) which indicates that the behavioral categorization of ambiguous face-hand morphs is related to the differential neural responses between face- and body-selective regions.

**Figure 5.**
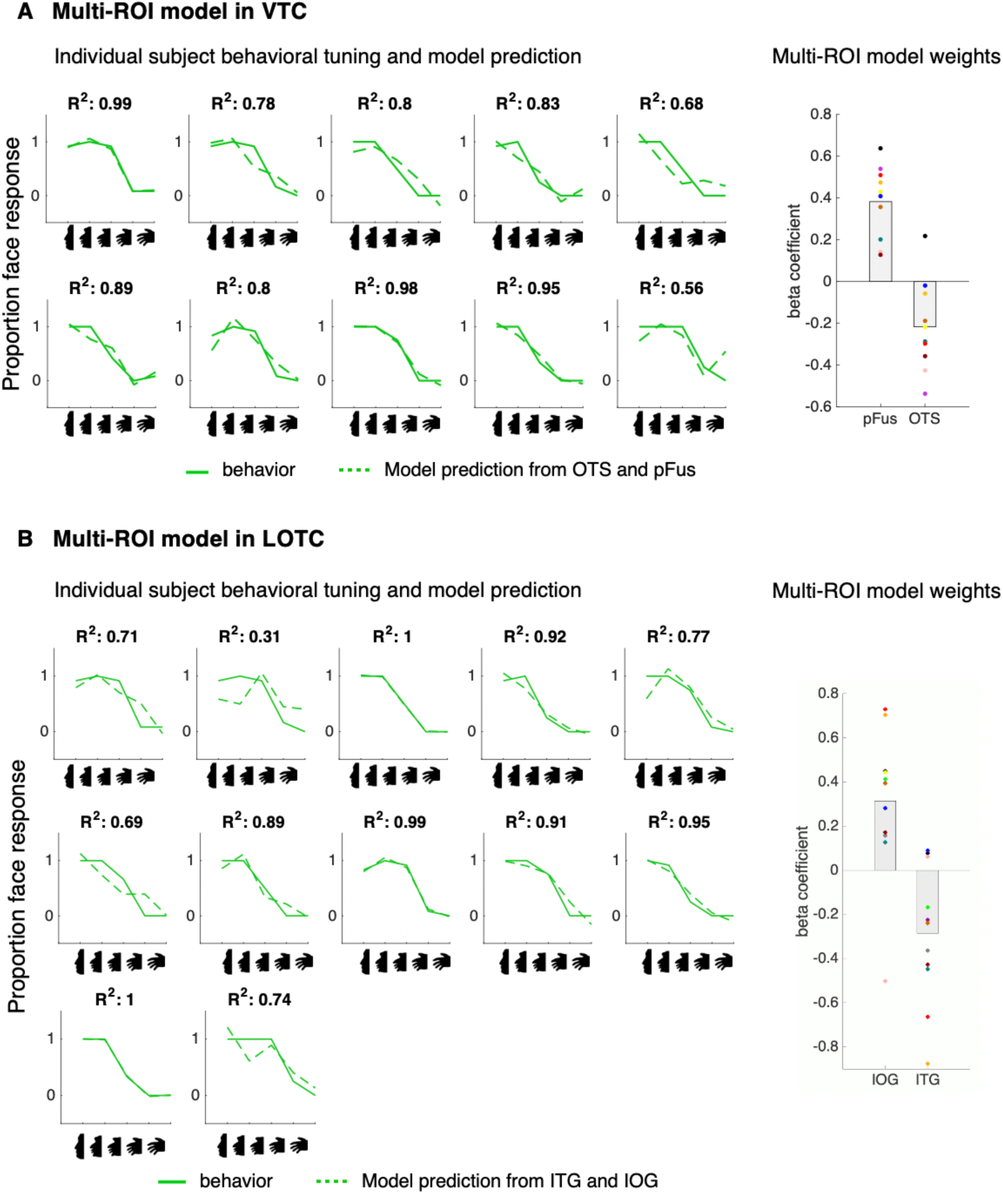
A multi-ROI model in VTC and LOTC successfully predicts behavioral performance. **A.** Left: Each panel shows the behavioral tuning (solid) for a participant and the MRM prediction using the neural tuning from both pFus-faces and OTS-bodies (pFus+OTS) in the right hemisphere (dashed). *X-axis:* morph level; *Y-axis:* proportion face response. R^2^: explained variance for each respective participant, indicated at the top of each subplot. **Right:** Beta coefficients of the right pFus+OTS MRM. The model assigned positive weights to pFus-faces and negative weights to OTS-bodies for most participants. *X-axis:* predictors; *Y-axis:* beta coefficients. Bars indicate the mean coefficient, and each dot indicates a participant. **B.** Identical analysis as in A but for the LOTC MRM using IOG-faces and ITG-bodies. Colors are the same as in prior figures.

### The MRM built from a group of participants accurately predicts behavioral tuning in a new participant

We reasoned that if the relationship between neural and behavioral tuning revealed by the MRM is a subject-general relationship between brain and behavior, then a model built from neural responses in one group of participants should be capable of predicting behavioral tuning in a new person. However, if the relationship between neural and behavioral tuning is subject-specific, such a model would not predict the behavioral tuning of a new participant very well. To investigate this possibility, instead of building an MRM for each subject, we built 1) a VTC MRM that relates behavioral categorization to neural responses from pFus-faces and OTS-bodies in N-1 participants, as well as 2) a LOTC MRM that relates behavioral categorization to neural responses from IOG-faces and ITG-bodies in N-1 participants. The outputs of each model are two coefficients of the linear contribution of the face and body ROIs (VTC: pFus-faces and OTS-bodies; LOTC: IOG-faces and ITG-bodies) to behavioral categorization (Fig 6A). We then used the model coefficients to predict behavioral tuning in a new subject based on the new subject’s neural tuning from either VTC (pFus-faces and OTS-bodies) or LOTC (IOG-faces and ITG-bodies), respectively. This approach was repeated using a 10-fold cross-validation procedure, using every subject once as a left-out test subject (Fig. 6B).

**Figure 6.**
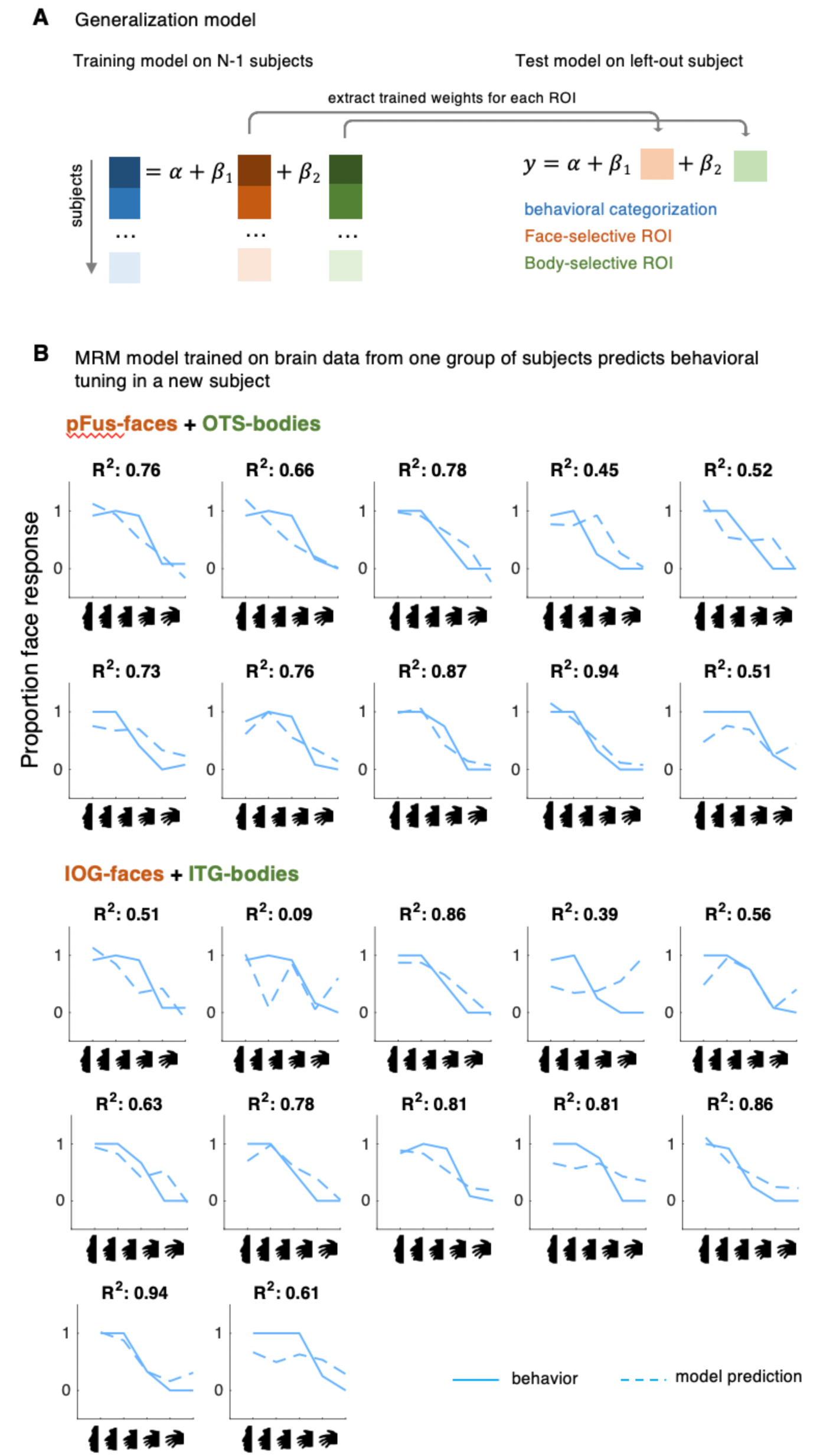
A cross-validated, multi-ROI model that is trained on brain data from one group of subjects predicts behavioral tuning in new subjects. **A.** Schematic illustration of the cross-validation procedure. *Left:* N-1 subjects were used to estimate model parameters relating behavioral categorization to neural tuning from a face-selective ROI and a body-selective ROI in the right hemisphere (see text for details). *Right:* The resulting beta coefficients were then multiplied with the fMRI data of the left-out-participant and were used to predict behavioral tuning for each morph level. B. In separate subplots, behavioral tuning (solid) for a participant and the model’s prediction of the right hemisphere MRMs (dashed) are shown. *X-axis:* morph level; *Y-axis:* proportion face response. *Solid blue line:* Behavioral tuning from the left-out participant. *Dashed blue line:* Model. R^2^: explained variance for each left-out participant is indicated at the top of each subplot. *Top*: VTC MRM. *Bottom*: LOTC MRM.

Impressively, the cross-validated VTC MRM using right hemisphere data accurately predicted behavioral tuning of the left-out subject (mean R^2^ ± SE: 0.70 ± 0.05, Supplemental Fig. 5). In contrast, predicting behavior from the VTC MRM based on left hemisphere data (mean R^2^ ± SE: 0.50 ± 0.09) was significantly worse (paired permutation testing, p < 0.01). Interestingly, the high predictability of the right hemisphere VTC MRM generalized to the cross-validated LOTC MRM (mean R^2^ ± SE: 0.66 ± 0.07), which performed equally well as the VTC MRM (permutation testing: p = 0.31, Fig. 6). Unlike the VTC MRM, there was no difference in performance of the LOTC MRM between hemispheres (permutation testing, p= 0.57). Finally, both cross-validated VTC and LOTC MRM models out-performed a right hemisphere cross-validated MRM using neural tuning from VO-2+CoS (mean R^2^ ± SE: 0.23 ± 0.08; both VTC and LOTC to VO-2+CoS model comparisons p<0.01), but only the LOTC MRM outperformed the VO-2+CoS control MRM in the left hemisphere (VO-2+CoS mean R^2^ ± SE: 0.26 ± 0.06; permutation testing vs. VTC MRM: p = 0.07, vs. LOTC MRM: p < 0.05).

Investigating the right hemisphere cross-validated VTC MRM more closely shows that the model captures the trend of each subject’s behavioral tuning even though the estimated beta coefficients were generated using neural tuning from an independent group of participants (Fig. 6B). Consistent with the within-subject MRM, the cross-validated VTC MRM displayed opposing weights for pFus-faces and OTS-bodies in that positive coefficients were derived for pFus-faces (β = 0.31±0.003) and negative coefficients for OTS-bodies (β = -0.22±.004). Likewise, the cross-validated LOTC MRM displayed opposing weights for IOG-faces (β = 0.20±.004) and ITG-bodies (β = -0.34±.004). While the cross-validated right hemisphere VTC MRM predicted behavior well overall, it is worth emphasizing that behavioral tuning deviated most from the neural predictions at the middle face-hand morph, which is the morph-level that was associated with the largest within-subject variability (Fig. 3). Further, participants that showed a lower explained variance in the within-subject MRM (Fig. 5B), also showed a lower explained variance in the cross-validated MRM (Fig. 6B). While the LOTC MRM, on average, explained a large amount of variance in both hemispheres, model fits deviate from the actual behavioral data to a greater degree compared to the VTC MRM (Supplemental Fig. 6, Fig.6B).

Altogether, these analyses reveal that (a) the combination of neural tuning from separate combinations of face- and body-selective regions in VTC and LOTC predict behavioral tuning to face-hand morphs, and (b) this relationship is subject-general. That is, behavioral tuning in a new participant can be predicted from a differential weighting of neural responses in face- and body-selective ROIs derived from a separate group of participants.

## DISCUSSION

In the present study, we measured neural tuning to face-hand silhouettes from face- and body-selective regions in human ventral temporal cortex (VTC) while participants viewed carefully controlled visual stimuli in the fMRI scanner. The visual stimuli were images that were morphed in a continuous shape space in which one side was a face silhouette, and the other a hand silhouette. The center shape of the continuum was perceived as being approximately equal parts of both categories. We then conducted a behavioral experiment in the same participants outside the fMRI scanner to examine the relationship between neural and behavioral tuning. Our results revealed that neural tuning from adjacent face- and body-selective regions together better predicted behavioral tuning compared to neural tuning from either type of region by itself. Furthermore, our results are not specific to VTC, but also extend to cortically adjacent face- and body-selective regions located in lateral occipito-temporal cortex (LOTC). Moreover, the relationship between neural tuning (in both VTC and LOTC) and behavioral tuning was consistent such that a model trained on neural tuning from one group of participants accurately predicted a new participant’s behavioral tuning.

Here, we propose that cortical adjacency likely enables the integration of neural signals between functionally-distinct regions. We then discuss how the best combination of neural tuning that predicts behavioral categorization may depend on the stimulus and task, as well as elaborate on the limitations of our findings. Finally, we discuss how the model-based approach implemented here is generalizable to future studies examining the relationship between neural and behavioral categorization.

### Cortical adjacency likely enables the integration of neural signals between distinct functional regions

Given that a large body of work in humans and non-human primates supports the causal role of domain-specific regions within high-level visual cortex in the perception of their preferred domain, the present findings showing that the combination of responses from pairs of face- and body-selective regions in VTC and LOTC best predict the categorization of face-hand morphs may seem surprising. However, when considering the fact that pFus-faces and OTS-bodies in VTC and IOG-faces and ITG-bodies in LOTC are two pairs of regions located within larger cortical expanses specialized for processing animate categories, the present findings are more intuitive. Indeed, our findings suggest that because face- and body-selective regions are cortically adjacent within a larger animate representation, it may be particularly beneficial for discriminating animate categories from one another in ambiguous situations. For example, cortical adjacency likely enables the combination of signals between regions through short-range connections and the combination of these neural signals likely contributes to categorical judgments.

More broadly, we propose that cortical adjacency is an implementational feature of the brain that enables the integration of information across domains, which is consistent with a sparsely-distributed organization of high-level visual cortex (Weiner and Grill-Spector 2010). Specifically, over the last twenty years, many researchers who use neuroimaging techniques to study the neural bases of object recognition have debated whether modular or distributed neural codes are more optimal for perceptual categorization (Baker et al. 2007; Op de Beeck, Haushofer, and Kanwisher 2008; Charest et al. 2014; Cichy et al. 2019; P E Downing et al. 2001; Golarai et al. 2007; Kriegeskorte et al. 2008; Lindh et al. 2019; Mur, Bandettini, and Kriegeskorte 2009; Tarr and Gauthier 2000; Weiner and Grill-Spector 2015).

Prior to the advent of fMRI, a long history of anatomy research stressed that both modular and distributed components together may be optimal for different types of information processing: segregation and integration, respectively. In terms of segregation and modularity, parallel processing could occur in separate functional regions, which would increase the speed and efficiency of information processing. Additionally, the integration of distributed information between regions would be beneficial to generate higher representational capacity as well as flexible processing of visual inputs (Felleman and Van Essen 1991; Grill-Spector and Weiner 2014; Zeki and Shipp 1988). We recently applied these classic anatomical concepts to explain the functional organization of human VTC in which we characterized its organization as *sparsely-distributed*. Sparseness refers to the presence of a series of minimally overlapping, highly-selective clusters that are arranged in a consistent topography relative to one another as well as retinotopic visual areas, while distributed refers to the fact that despite the minimal overlap between clusters, there is a substantial amount of category information across voxels with different functional properties. Our present data indicate that cortical adjacency may be a fine-scale implementational feature supporting this sparsely-distributed organization. As discussed previously (Grill-Spector and Weiner 2014), the segregation of face- and body-selective regions into distinct domain-specific networks allows parallel processing of category information in non-ambiguous situations, and is consistent with theories of modularity (N. Kanwisher 2010). Additionally, the adjacency of face- and body-selective regions enables cross-talk and integration of information across the domains of faces and bodies – for example, linking a person’s face and body, or perception under ambiguity, which are both consistent with distributed theories of VTC (J V Haxby et al. 2001). Thus, cortical adjacency likely functions to maximize computational benefits from both sparse (localized cortical clusters in functionally specialized networks) and distributed (the integration of neural signals between networks) organizational features of high-level visual cortex.

Consistent with our results and the interpretation of our findings, recent anatomical and functional connectivity studies in macaques also propose the combination of neural signals between category-specific networks selective for faces and bodies. Indeed, while recent research in macaques identifies distinct cortical networks that separately process face and body information (Borra et al. 2008; Moeller, Freiwald, and Tsao 2008; Orban, Van Essen, and Vanduffel 2004; Pinsk et al. 2009; Tsao and Livingstone 2008), these studies also identify connections that are shared between these two networks (Borra et al. 2008; Premereur et al. 2016). These studies propose that the subset of these networks that overlap with one another may be integral for the combination of face and body information both anatomically (Borra et al. 2008) and functionally (Premereur et al. 2016). Thus, future studies examining the relationship between neural tuning and behavioral categorization while recording responses from single neurons within adjacent face- and body-selective regions can test if this combination occurs at the neuronal level.

### The best combination of neural tuning that predicts behavioral categorization may depend on the stimulus and task

Our present findings revealed that neural tuning from certain combinations of face- and body-selective regions more accurately predicted behavior than other combinations of neural tuning of face- and body-selective regions. For example, in VTC, the combination of neural tuning from pFus-faces and OTS-bodies better predicted behavioral tuning compared to the combination of neural tuning from mFus-faces and OTS-bodies – especially in the right hemisphere. We hypothesize that the improved prediction of behavioral categorization from pFus-faces and OTS- bodies rather than mFus-faces and OTS-bodies may be related to the stimuli and the respective location of pFus-faces and mFus-faces in the visual processing hierarchy. As mFus-faces is situated at a higher position in the visual hierarchy compared to pFus-faces (Grill-spector et al. 2017; Grill-Spector et al. 2018), we hypothesize that the present stimuli (simple, two-dimensional shapes) were more suitable for pFus-faces. This hypothesis is also consistent with our LOTC findings, as IOG-faces is considered to be at an even earlier position in the visual hierarchy compared to pFus-faces (Grill-Spector et al. 2017, 2018; James V. Haxby, Hoffman, and Gobbini 2000; Rossion 2008). We also propose that behavioral categorization of more complex stimuli may be attained from the linear combination of neural tuning from mFus-faces and OTS-bodies than the combination of either pFus-faces and OTS-bodies or IOG-faces and ITG-bodies, which can be tested in future research. In addition, future research should assess on a trial-by-trial basis how neural signals predict single images in real time and with a 1:1 mapping between behavioral categorization and neural responses.

Future studies can also examine how combinations of neural tuning affect the lateralization effects identified in VTC. For example, recent proposals hypothesize that the lateralization for word processing in the left hemisphere and face processing in the right hemisphere develops from the competition of foveal resources during development in combination with the lateralization of the language system in the left hemisphere (Behrmann and Plaut 2015; Dehaene et al. 2015). As face- and word-selective regions form a cortical cluster in left posterior VTC, while face-, body-, and word-selective regions form a cluster in left mid-VTC, future studies examining the relationship between neural tuning and behavioral tuning during categorization tasks that involve faces, bodies, and words may find that combinatorial tuning from all three regions in the left, but not the right, hemisphere would out-perform other models. Altogether, we propose that the cortical adjacency of functional regions in VTC likely enables flexible combinations of neural tuning that can accommodate different types of stimuli and task demands.

### Limitations

Using simplified stimuli to investigate complicated experimental questions such as those related to the perception of visual ambiguity is beneficial because the images are generated on a known continuum of shape space in which (a) the ends of the continuum represent shapes from two different categories and (b) there is a clear categorical transition such that the center shape of the continuum is composed of equal parts of both categories. Such a space enables a clear quantification of how neural and behavioral responses change when exposed to stimuli that cross category boundaries.

Despite the controlled feature space of these stimuli, a concern may be that the brain-behavior relationships modeled in the present study may not generalize to naturalistic stimuli (Nishimoto et al. 2011; Huth et al. 2012, 2016; Naselaris et al. 2012). We acknowledge this limitation and propose that our present results and model form a foundation from which to build more complex brain-behavior models (Rust and Movshon 2005) underlying the perception of visual ambiguity – especially for animate categories as silhouettes of animate objects are more accurately recognized than other types of silhouettes (Lloyd-Jones and Luckhurst 2002; Tian et al. 2016). Nonetheless, we underscore that neurally, silhouettes are sufficient to selectively drive high-level category-selective regions within VTC, as face- and body-selective regions respond selectively to silhouettes of their preferred category, shown in the present study as well as in previous work (Andrews et al. 2002; Stefania Bracci et al. 2010; Davidenko, Remus, and Grill-Spector 2012; Hasson et al. 2001; Mendola et al. 2018; Tong and Nakayama 1999). We also emphasize that, to our knowledge, these data are the first to show that regions from different domain-specific networks in VTC and LOTC can work together to achieve a behavioral goal. Future studies can extend our work and test additional categories as well as formats in order to examine if a) the combination of neural responses between functional regions best predicts behavioral responses to visually ambiguous stimuli and b) these brain-behavioral predictions are indeed more prominent for animate categories.

### Beyond the representation of animate categories in VTC

The linear model-based approach we have implemented in the present study is applicable to additional factors beyond the representation of animate categories in VTC. For example, neurally, previous studies have examined visual categorization in brain areas outside of VTC (Freedman et al. 2001; Jiang et al. 2007; Ferrera et al. 2009), as well as have considered how other brain areas (for example, within prefrontal cortex (Baldauf and Desimone 2014)) may act as gatekeepers that guide the categorical decision process within cortical regions in VTC. Additionally, categorization is not limited to the visual domain. For instance, empirical studies examine the categorization of (a) movements in motor cortex (Seger and Miller 2010), (b) odors in piriform cortex (Tantirigama et al. 2017), and (c) sounds in the visual cortex of blind individuals (van den Hurk et al. 2017). The approach we have implemented here can be applied to these examples, as long as two requirements are fulfilled. First, there should be a stimulus continuum between (at least) two categories. Second, there must be a hypothesized relationship between behavioral tuning and neural tuning from the functional regions of interest. As long as these two requirements are met, our model-based approach can be applied to additional types of categorization, as well as additional functional regions in future studies. Further, we also emphasize that while we establish this approach with neural tuning that is read-out from entire functional regions, the spatial scale of this approach is presently unknown. For instance, distinct cortical columns in different parts of the brain represent (a) different viewpoints of faces (Tanaka 1996), (b) distinct motion directions (Albright 1984), or (c) specific parts of the body surface (Mountcastle 1997). Future studies can test if the combination of neural tuning signals from multiple cortical columns better predicts behavioral categorization compared to neural tuning signals from single cortical columns alone.

### Conclusions

In summary, we find that neural categorization tuning from multiple category-specific regions in human ventral temporal and lateral occipito-temporal cortices accurately models behavioral categorization tuning. Furthermore, this model trained on the neural tuning from one group of participants nicely aligns wFsuith behavioral tuning in new participants with a right hemisphere dominance. Together, these data suggest that adjacent cortical regions with opposed neural tuning can accommodate efficient and flexible categorization.

## Acknowledgements

We thank Kendrick Kay for useful discussions regarding statistical analyses. This research was supported by UC Berkeley start-up funds (to KSW) and NEI grant: 1R01EY02391501A1 (to KGS).

## Supplemental Figures

**Supplemental Figure 1.**
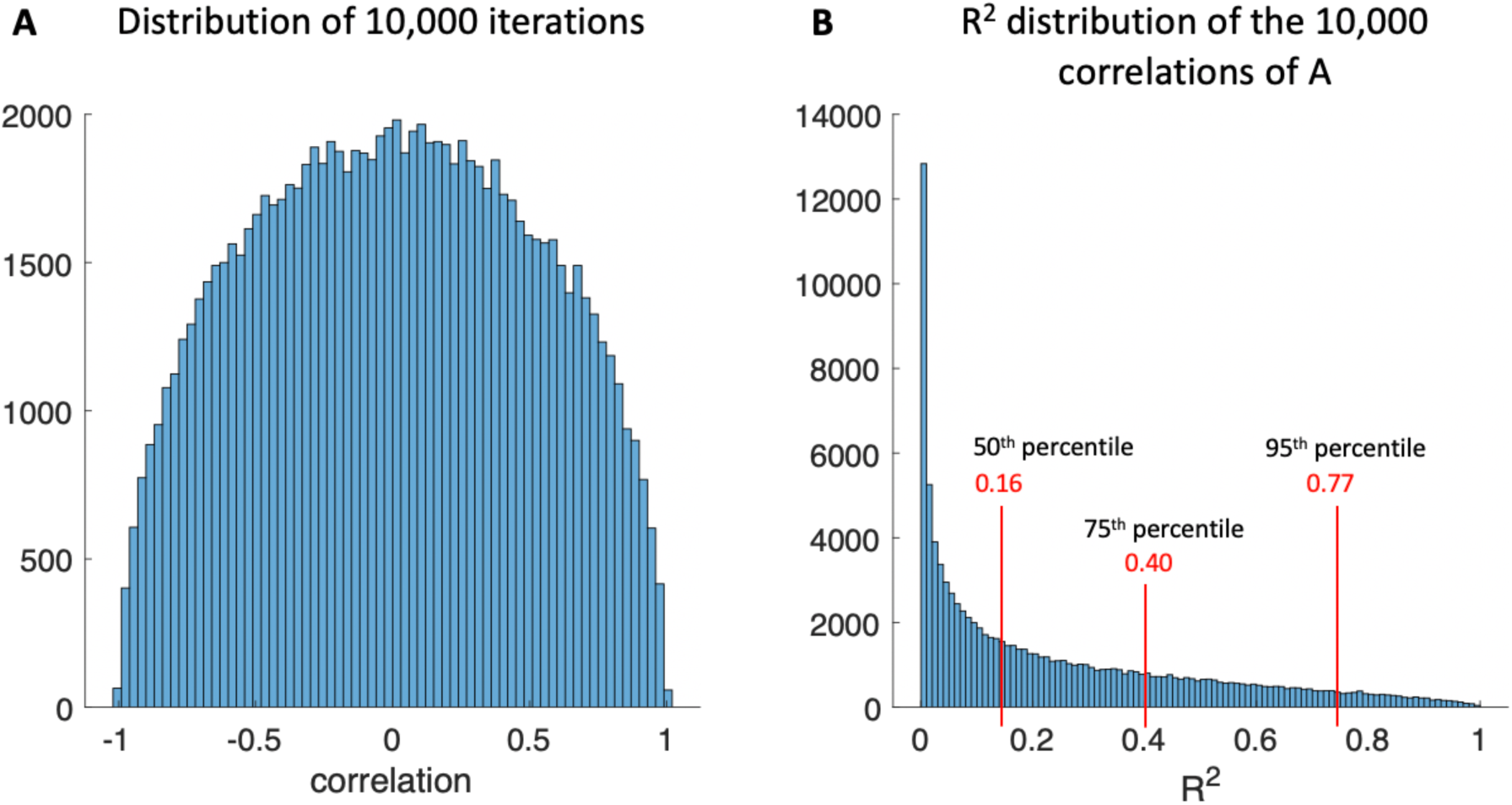
Simulation of 10,00 5-point correlations. **A.** In order to establish a random null distribution of of correlations of 5 points that are similar to our neural responses and behavioral data, we generated 10,000 random 5-point vectors stemming from (1) a normal distribution, comparable to our z-scored beta values of one ROI, and (2) a vector of 5 random numbers from a uniform distribution, comparable to our behavioral categorization responses.X-axis: Correlation values of two vectors, y-axis = frequency of correlation occurrence. **B.** R^2^ distribution of the 10,000 correlations established in A. R^2^ was calculated by squaring the correlation value. Red lines indicate the 50^th^, 75^th^ and 95^th^ percentile, respectively. X-axis = explained variance (R^2^), y-axis = frequency of that R^2^ occurance.

**Supplemental Figure 2.**
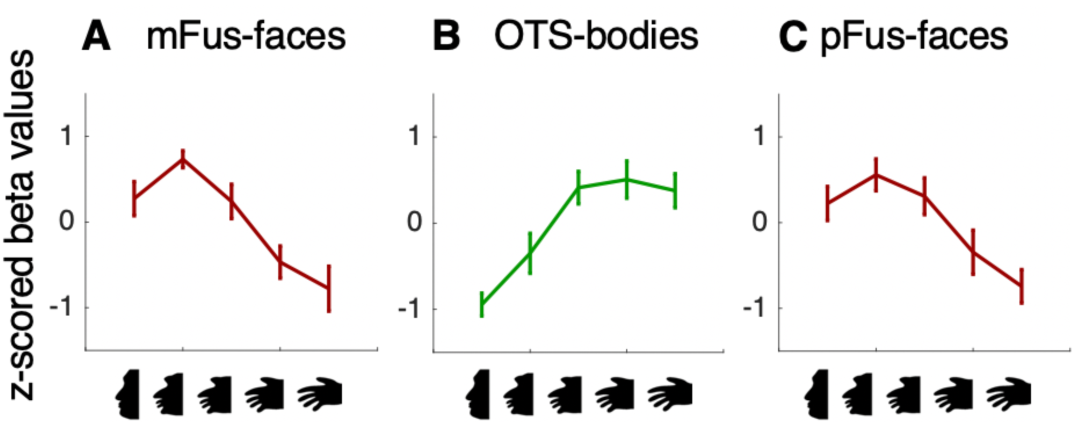
Face- and limb-selective regions in left ventral temporal cortex (VTC) display differential neural tuning. Same layout as Fig. 2, but for the left hemisphere. Beta responses to each of the 5 morph levels of face-hand silhouettes were extracted from each region in the left hemisphere and transformed into z-scores to generate normalized neural tuning for each participant. The mean neural tuning across participants is shown for each region of the left hemisphere, across participants. Face- and limb-selective regions have different neural tuning. *X-axis*: face-hand silhouette morph level. *Y-axis:* Z-scored beta values. *Errorbars:* Standard error across subjects.

**Supplemental Figure 3.**
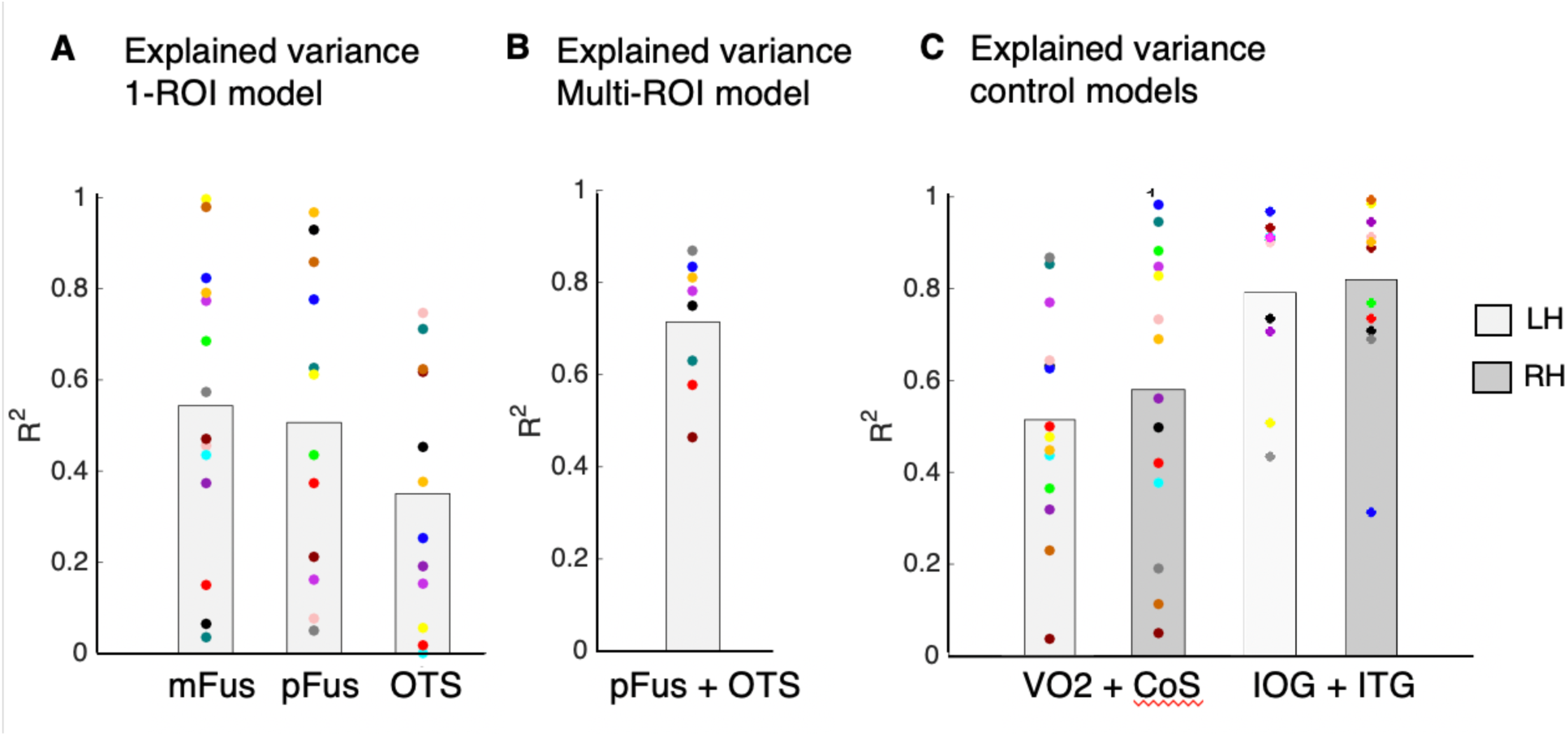
Establishing the relationship between neural tuning and behavioral tuning. A and B: Same layout as Figs. 4C and D, but for the left hemisphere. **A.** Explained variance (R^2^) from the 1-ROI model in the left hemisphere. Neural tuning from each ROI was a significant predictor of behavioral tuning. *X-axis:* ROIs used as predictors for the linear regression model. Variance explained of the behavioral data by three 1-ROI models. Bars indicate the mean across participants, and each dot represents data from one participant. The color of each participant is matched across models and data in all other figures. **B.** Variance explained of the behavioral data by a multiple ROI model (MRM) including left pFus-faces and left OTS-bodies. Conventions same as A. **C.** Explained variance of the control models using retinotopic area VO-2 and place-selective region CoS-places as well as IOG-faces and ITG-bodies. Conventions same as in A and B. Individual subject dots are color matched. Light-gray bars: Mean across participants for the left hemisphere. Dark-gray bars: Mean across participants for the right hemisphere. LH = left hemisphere, RH = right hemisphere.

**Supplemental Figure 4.**
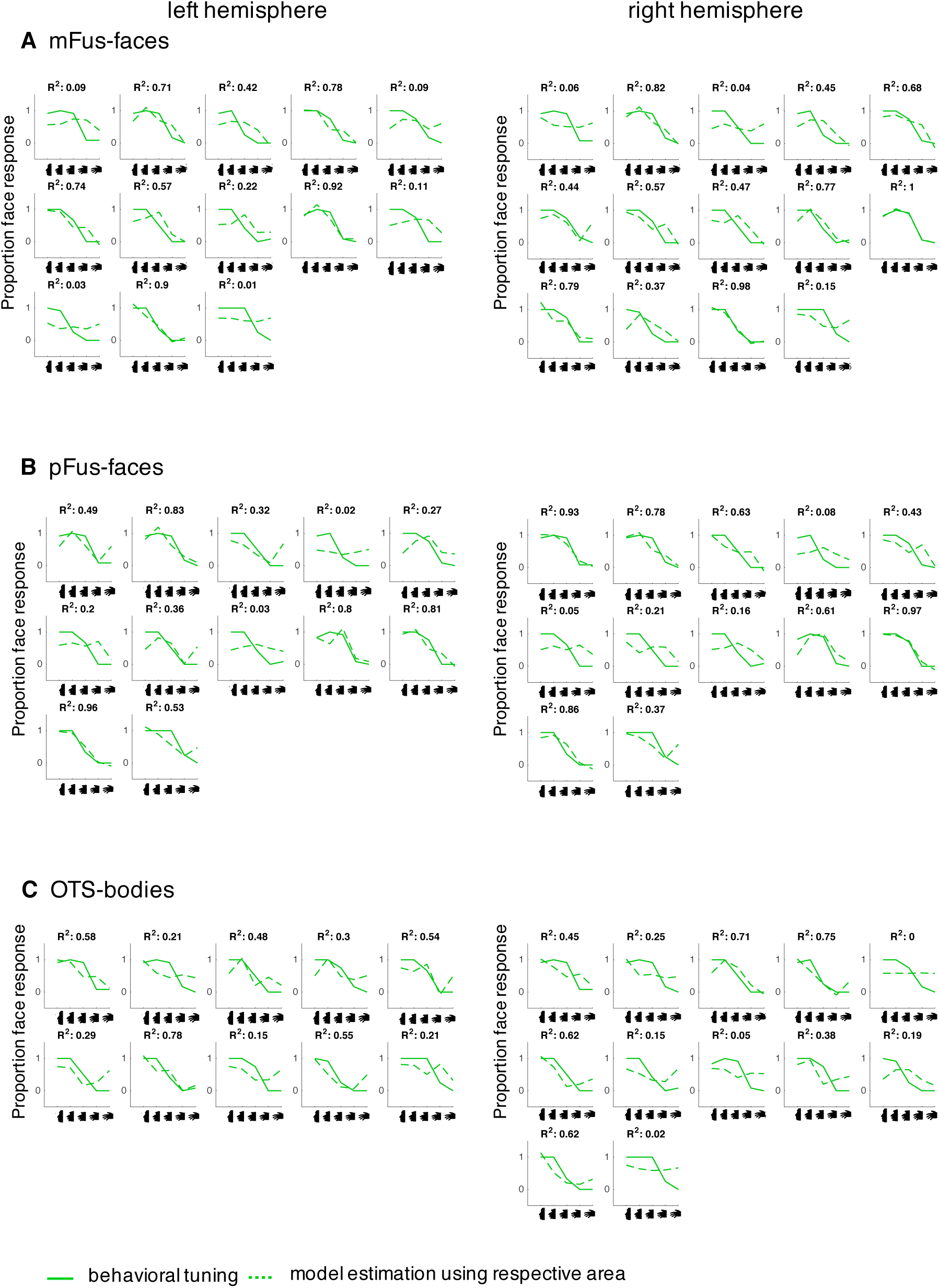
Model-generated behavior vs. actual behavior in individual participants of the 1-ROI models. **A-C.** Behavioral tuning (solid) for each participant and model estimation (dotted) of behavioral tuning from the linear regression model using mFus-faces (A), pFus-faces (B), and OTS-bodies (C). Each subplot corresponds to one participant. The number of subjects differs per plot since not every ROI could be defined within every subject (Supplmental Table 1). *X-axis:* morph level. *Y-axis:* proportion face response. Explained variance for the respective participant is shown at the top of each subplot. *Left column:* Left hemisphere data. *Right column:* Right hemisphere data.

**Supplemental Figure 5.**
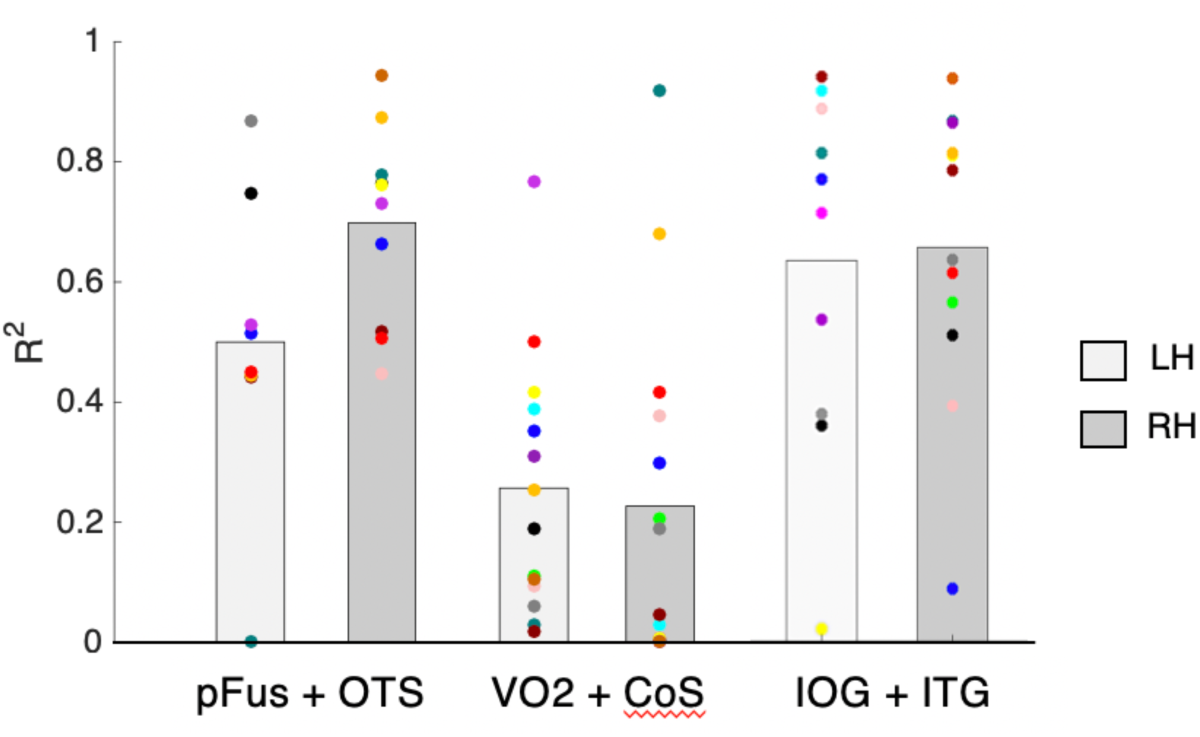
Explained variance of the cross-validated MRM models across participants. Explained variance (R^2^) of the cross-validated MRM models. N-1 subjects were used to determine coefficient weights for pFus-faces and OTS-bodies. The resulting beta coefficients were then multiplied with the fMRI data of the left-out-subject and used to predict their behavioral tuning for each morph level. *X-axis:* ROIs used as predictors for the linear regression model. *Y-axis:* Bars indicate the mean across participants, and each dot represents data from one participant. The color of each participant is matched across models and data in all other figures. Light-gray bars: Mean across participants for the left hemisphere. Dark-gray bars: Mean across participants for the right hemisphere.

**Supplemental Figure 6.**
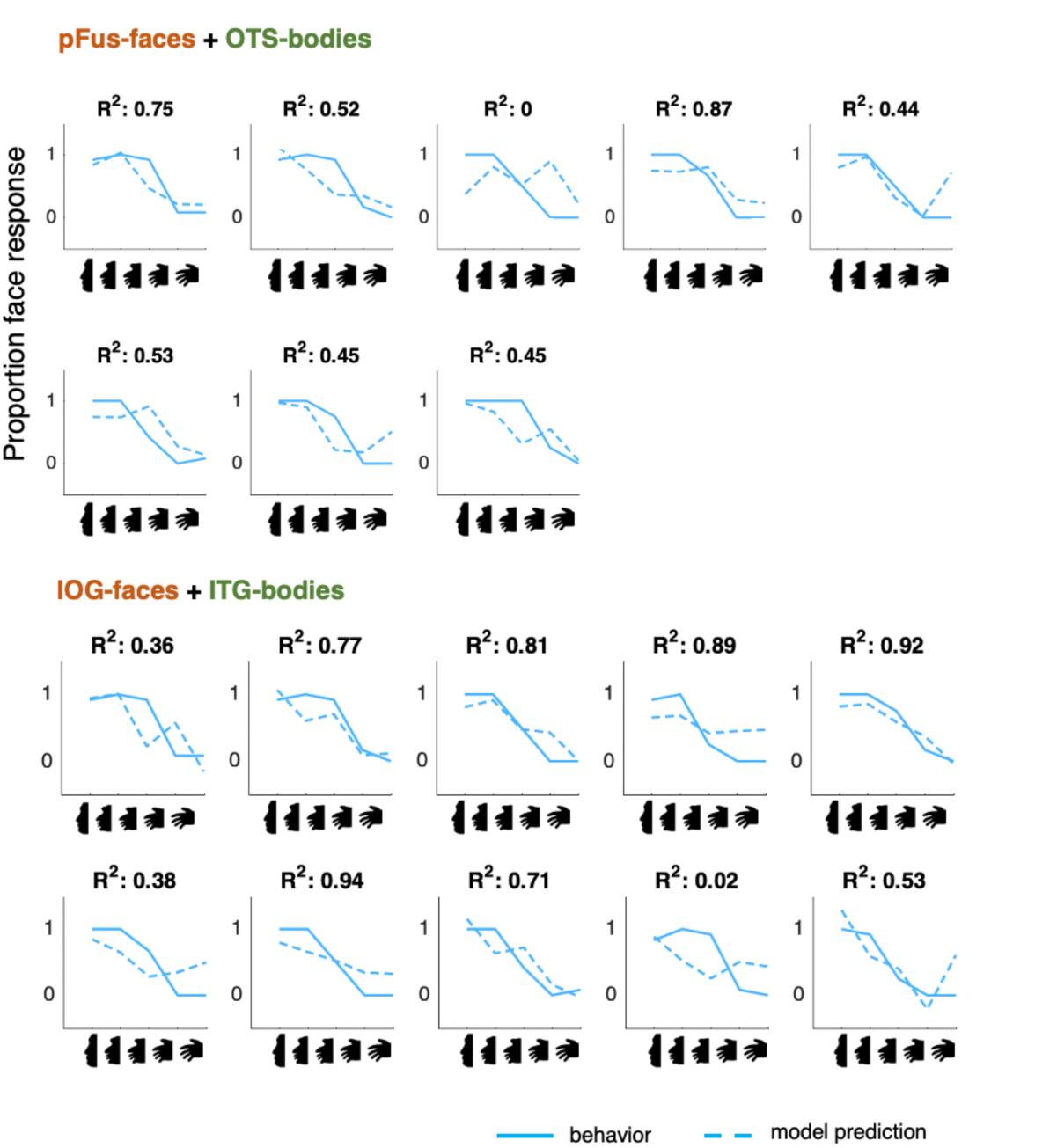
A cross-validated, multi-ROI model that is trained on brain data from one group of subjects predicts behavioral tuning in new subjects. In separate subplots, behavioral tuning (solid) for a participant and the model’s prediction of the *left hemisphere* MRMs (dashed) are shown. *X-axis:* morph level; *Y-axis:* proportion face response. *Solid blue line:* Behavioral tuning from the left-out participant. *Dashed blue line:* Model. R^2^: explained variance for each left-out participant is indicated at the top of each subplot. Top section: VTC Multi-ROI model, Bottom section: LOTC Multi-ROI model.

## Supplemental Tables

**Supplemental Table 1.**
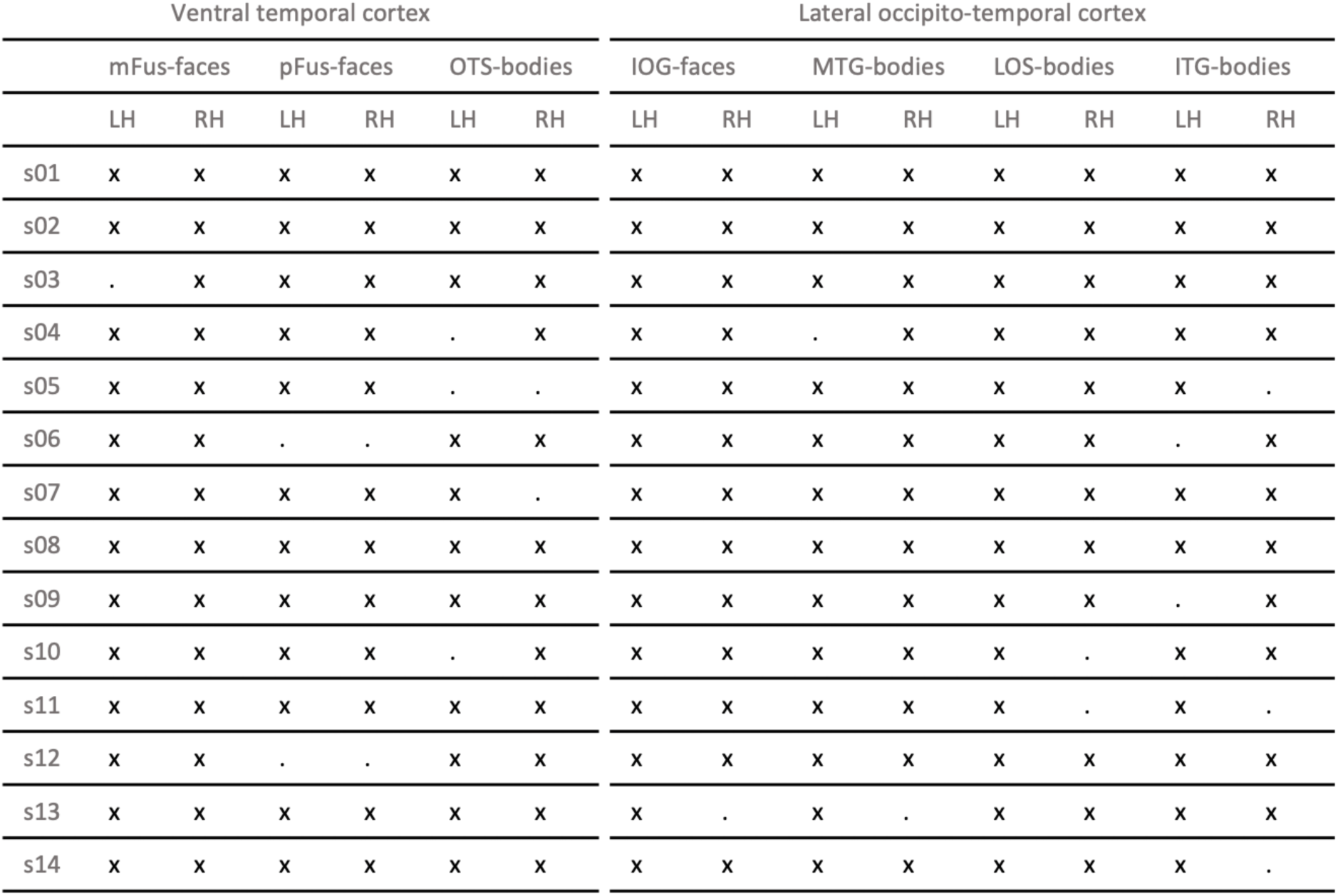
Frequency of face-selective and body-selective regions identifiable in each subject within VTC and LOTC. For each subject (rows), we report whether we were able to define the given ROI using a common statistical threshold (*t* > 3, voxel-level). Abbreviations: *LH:* Left hemisphere, *RH:* right hemisphere, *x* = ROI defined, . = no ROI defined.

**Supplemental Table 2.**
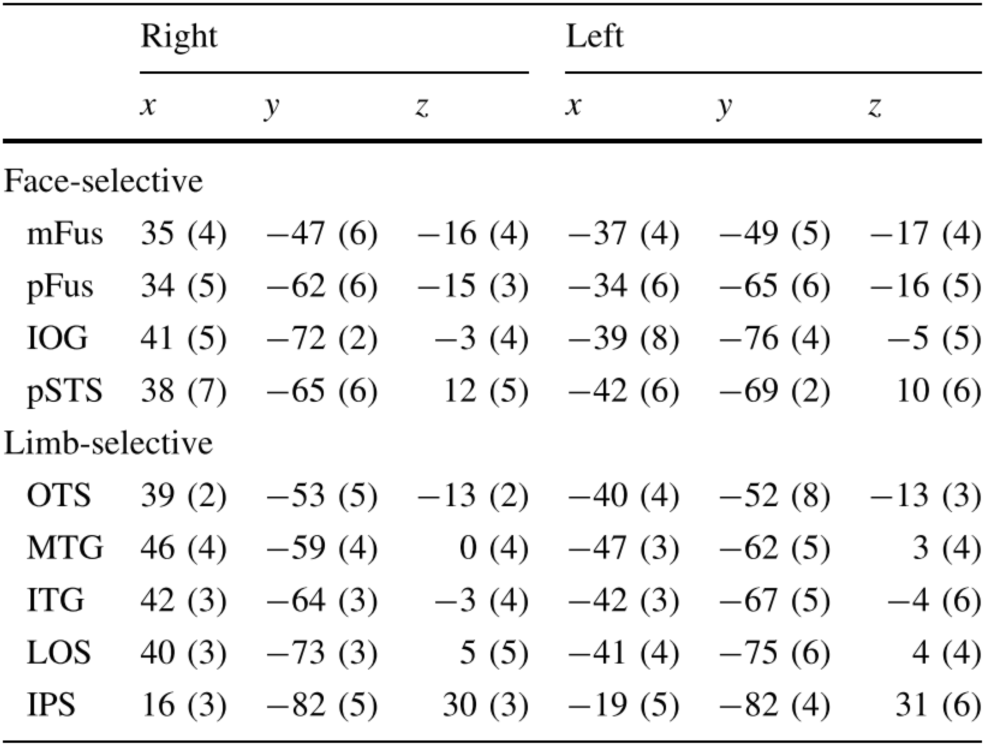
Location of face- and limb-selective regions in Talairach space, Table 1 from Weiner and Grill-Spector (2013). Standard deviations are across 9-11 subjects.

